# Logarithmic coding leads to adaptive stabilization in the presence of sensorimotor delays

**DOI:** 10.1101/2025.04.28.650940

**Authors:** Leonardo Demarchi, Monica Coraggioso, Antoine Hubert, Thomas Panier, Ghislaine Morvan-Dubois, Volker Bormuth, Georges Debrégeas

## Abstract

Animals respond to sensory stimuli with motor actions, which in turn generate new sensory inputs. This sensorimotor loop is constrained by time delays that impose a trade-off between responsiveness and stability. Additionally, as the relationship between a motor command and the corresponding sensory feedback is context-dependent, the response must be adapted in real time. It is generally believed that this adaptation process relies on an internal model that is continuously updated through prediction error minimization. Here, we experimentally reveal an alternative strategy based on a simpler feedback mechanism that does not require any internal model. We developed a virtual reality system for the miniature transparent fish *Danionella cerebrum* that enables in vivo brain-wide imaging during fictive navigation. By systematically manipulating the feedback parameters, we dissected the motor control process that allows the animal to stabilize its position using optic flow. The sensorimotor loop can be fully described by a single delay differential equation, whose solutions quantitatively capture the observed behavior across all experimental conditions. Both behavioral and neural data indicate that the observed adaptive response arises from the logarithmic nonlinearities at the sensory (Weber-Fechner law) and motor (Henneman’s size principle) ends. These fundamental properties of the nervous system, conserved across species and sensory modalities, have traditionally been interpreted in terms of efficient coding. Our findings unveil a distinct functional role for such nonlinear transformations: ensuring stability in sensorimotor control despite inherent delays and sensory uncertainty.

---

Animals continuously interact with their environment through sensorimotor loops. These dynamic processes are inherently constrained by finite time delays and noise in the sensory and motor systems which can lead to instability [1, 2]. Moreover, the relationship between motor commands and the resulting sensory feedback is not fixed but rather depends on the state of both the body and the external world. For a given context, there exists an optimal response function that maximizes behavioral performance [3]. As the context varies, animals need to adapt their response in a flexible manner [4]. A key example is optic flow navigation, where animals use visual cues to estimate their motion in order to maintain an intended trajectory [5, 6]. This behavior is ubiquitous among motile species, but it is particularly important for flying and swimming animals, which can be easily thrown off course by external currents. The relationship between the animal speed and the retinal optic flow is ambiguous, as it depends on the distance of the animal from the surrounding objects, a parameter that varies in time and cannot be directly sensed by the animal. One way to solve this problem consists in creating and then constantly updating an internal model of the environment through a process of prediction-error minimization [7,8]. However, it remains unclear whether adaptive responses can emerge without such an internal model.

We address this question using the miniature freshwater fish *Danionella cerebrum*, a recently introduced model vertebrate whose size and transparency enables in-vivo whole-brain imaging across all developmental stages [9, 10]. We first use a virtual reality system to dissect the sensorimotor computation at play during optic flow navigation. We then use whole-brain imaging to probe the dynamics of the neuronal population engaged in this process.

## Results

### Stabilization of visual flows in a virtual reality system

The fish were head-tethered using agarose, then placed in a virtual-reality setup that comprises a camera to monitor tail movement, a projector to display visual patterns beneath the animal, and a light-sheet microscope for brain imaging (Fig. 1A, Methods). Experiments were performed on two-week-old danionella, which, unlike zebrafish larvae, exhibit continuous swimming in both freely moving and head-tethered configurations, with a quasi-constant tail-beat frequency of *≈* 16 Hz [11]. Using a fluid dynamics model calibrated against freely swimming recordings, we inferred the fictive forward speed *V* and angular velocity of the fish in real-time from the video-monitored tail movement [12, 13] (Fig. 1B, Methods). These values were used to compute the visual feedback, i.e. the translational and rotational velocity of the projected pattern to in-duce the illusion of self-motion in the fish.

**Figure 1:**
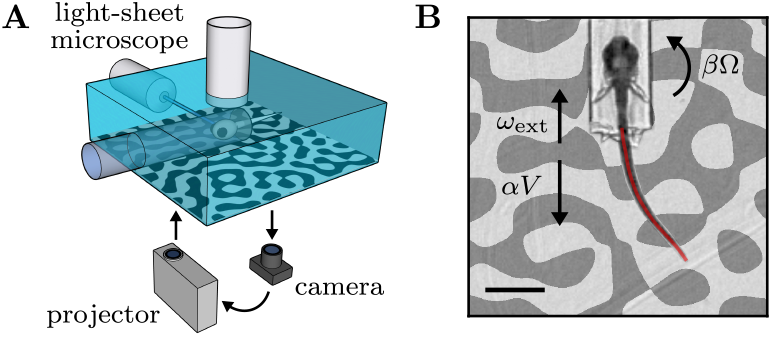
Virtual reality system. (A) Schematic of the experimental setup. Fish are head-tethered in agarose. Their tail movements are video-monitored and a projector displays a visual pattern on the opaque bottom of the tank. Brain-imaging can be simultaneously performed using a light-sheet microscope. (B) Snapshot of a fictively swimming fish. From successive tail profiles (red line), we extract the fictive speed *V*, and angular velocity *Ω* which are used to compute the translational and rotational feedback flow rates *αV* and *βΩ*, respectively. We additionally simulate an external current with flow rate *ω*_ext_. Scale bar: 1 mm.

In the real world, a movement of the fish at a certain speed corresponds, from the animal’s egocentric perspective, to an opposite movement of the environment with the same speed. However, what the animal actually perceives is the resulting optic flow on its retina, which only depends on the ratio between the speed and the distance of the surrounding objects. For a pattern moving at speed *v*, the optic flow is characterized by the rate *ω* = *v/h*, where *h* is the pattern distance, which in our case is 5 mm (Fig. 2A). We implemented the feedback by translating the pattern backwards, resulting in a flow rate *αV*, where the feedback gain *α* can be interpreted as the inverse of the pattern distance in the virtual environment. Note that the ambiguity in relating optic flow and self-motion is specific to translational movement: the angular velocity can be directly inferred from the rotational optic flow. We implemented rotational feedback by rotating the pattern with angular velocity *βΩ*, where a rotational feedback gain *β* = 1 corresponds to freely swimming conditions.

**Figure 2:**
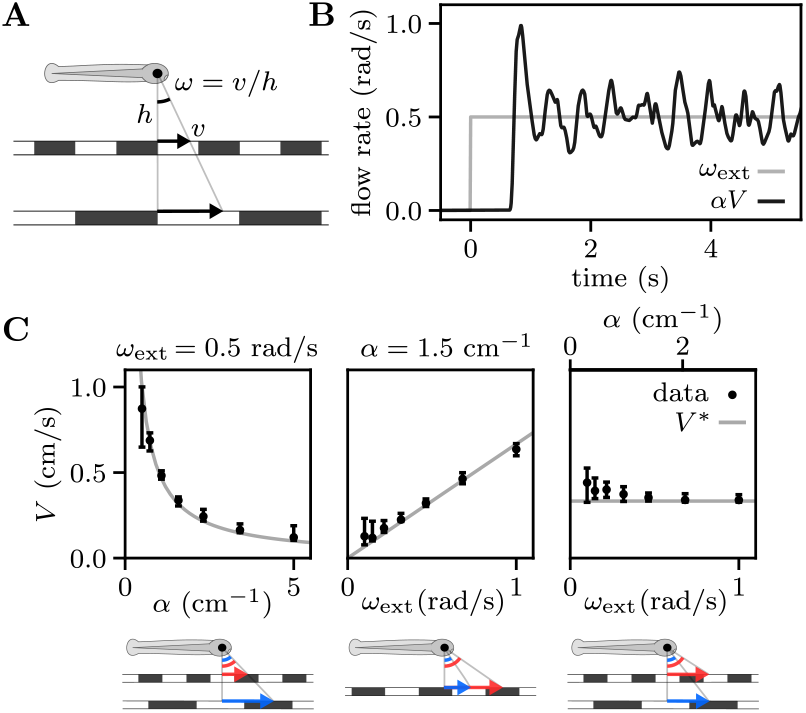
Fish adapt their swimming speed to stabilize external visual flows. (A) Definition of the optic flow rate *ω*. Rescaling the pattern size, distance *h* and speed *v* by the same factor does not affect the perceived optic flow. (B) Example trace showing the fish response to forward pattern motion. After a short latency, the fish adjusts its swimming speed to match external and feedback flow rates. (C) Median swimming speed (black error bars indicate quartiles of the distribution across *N* = (33, 31, 28) fish, from left to right) for different values of the external flow rate *ω*_ext_ and feedback gain *α*. Fish can adapt their swimming speed to the target value (gray lines) to stabilize themselves against currents. The schematics below the graphs illustrate the equivalence of the parameter changes in the real world, with the arrows denoting the speed of the external current for two different parameter values (larger in red).

We elicited an optomotor response by translating the pattern in a certain direction with an imposed rate *ω*_ext_, simulating an external current dragging the fish in the opposite direction [14]. The fish responded to such a stimulus by actively turning to align with the fictive current (movie S1). Here we focus on the case where the external current starts off directed along the fish heading direction, corresponding to a fictive backward movement. If the fish remains approximately aligned with the current then the optic flow is directed along the fish heading with net rate *ω* = *ω*_ext_ *− αV*.

The absence of visual feedback (*α* = 0, open-loop) rapidly induces the fish to give up and stop swimming. Otherwise (*α >* 0, closed-loop), they can keep swimming for sustained periods of time and adapt their swimming speed to match external and feedback flow rates such that *V ≃ V* ^*\**^ = *ω*_ext_*/α* (Fig. 2B, movies S2-3). Equivalently, this means that their position in the virtual space remains approximately constant.

We assessed the robustness of this behavior by presenting fish with external currents of varying optic flow rate *ω*_ext_ and for different values of the feedback gain *α*, in randomized trials lasting 30 s. We considered three experimental paradigms (Fig. 2C): fixed external flow rate (varying *α* with fixed *ω*_ext_), fixed pattern distance (varying *ω*_ext_ with fixed *α*), fixed current speed (varying *ω*_ext_ and *α* with a fixed target speed *V* ^*\**^). We found that most fish adapt their speed to compensate for the external current, provided that the target speed *V* ^*\**^ falls within a physiologically accessible range (SI section 1.7).

### Sensorimotor delays induce sustained speed oscillations

We noticed that the fictive speed signals displayed quasiperiodic oscillations around the target value *V* ^*\**^, as illustrated in Fig. 2B. These speed modulations have a characteristic frequency of *≈* 1.5 H and an amplitude proportional to the target speed (SI section 1.8-9). We hypothesized that they could be a signature of delays in the sensorimotor loop. Intuitively, if the correcting action lags the driving signal by a finite delay *τ*, the fish may systematically overcompensate for the same duration after crossing the target speed, leading to oscillations of period *∼* 4*τ*.

We measured the sensorimotor delay by modulating the imposed optic flow with a white-noise signal, while monitoring the fish velocity. The impulse response function, defined as the acceleration signal evoked by an impulse of optic flow rate, was then estimated by reverse correlation [15]. This response function exhibits a sharp delayed positive peak, corresponding to the response kernel, followed by a negative modulation reflecting the feedback mechanism (Fig. 3B).

**Figure 3:**
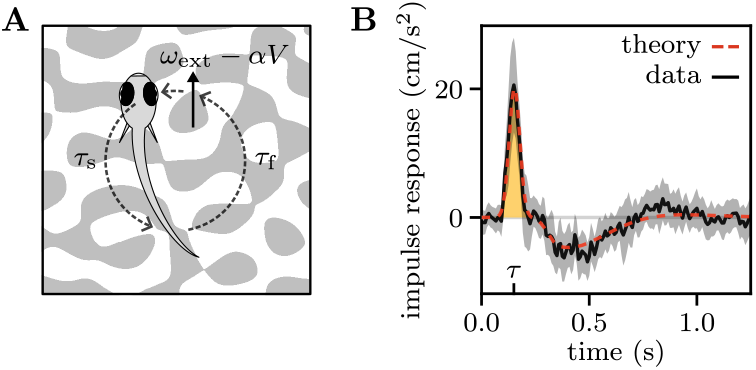
Time delays in the sensorimotor loop. (A) Schematic showing the delays in the sensorimotor loop. The fish perceives a certain optic flow and updates its swimming speed after a sensorimotor delay *τ*_*s*_. A given tail movement induces a change in the optic flow after a feedback delay *τ*_*f*_. (B) Reverse correlation gives an estimate of the impulse response function (black line and gray area, mean and standard deviation over *N* = 11 fish), i.e. the evolution of 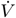 following an impulse perturbation of *ω*_ext_.The initial peak represents the response kernel (yellow area), whose time gives an estimate of the delay *τ*. Integrating Eq. 1 with this response kernel, we obtained the theoretical impulse response shown in red.

The time of the peak provides an estimate of the delay *τ* = 150 *±* 20 ms. This value is close to one-fourth of the oscillatory period observed in fish swimming under constant current flow (Fig. 2B), supporting our initial hypothesis. This sensorimotor delay can be decomposed into two components, *τ* = *τ*_s_+*τ*_f_, where *τ*_s_ corresponds to the internal delay between the sensory perception and the execution of the correcting motor action, while *τ*_f_ accounts for the lag between the tail motion and the resulting change in optic flow (Fig. 3A). While in freely swimming conditions *τ*_f_ is due to inertia, here it corresponds to the latency of the virtual reality, which we measured to be *τ*_f_ = 60 *±* 4 ms. Therefore, the sensory-to-motor internal delay is *τ*_s_ = 90 *±* 20 ms, of the order of a tail-beat period. This value was found to be independent of both *ω*_ext_ and *α* (SI section 2.2).

To model the sensorimotor loop, we first assumed that the fish acceleration at time *t* is proportional to the net flow rate at time *t − τ* :

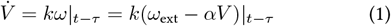

where the parameter *k* denotes the responsiveness of the fish to optic flow. Rewriting Eq. 1 in dimensionless form, we found that the system dynamics are governed by a single dimensionless gain, *µ* = *kατ*, whose value defines the stability regime (Fig. 4A, Methods). For *µ <* 1*/e, V* (*t*) decays exponentially to the target value *V* ^*\**^. For 1*/e < µ < π/*2, it converges to *V* ^*\**^ through damped oscillations. At *µ* = *π/*2, one observes a Hopf bifurcation, followed by an unstable regime with diverging oscillations. This equation could thus lead to the sustained oscillations observed in our data through two distinct mechanisms: in the intermediate regime, sensorimotor noise could drive the system away from the target speed eliciting noise-induced oscillations; in the unstable regime, nonlinearities could prevent the system from diverging leading to limit cycle oscillations.

**Figure 4:**
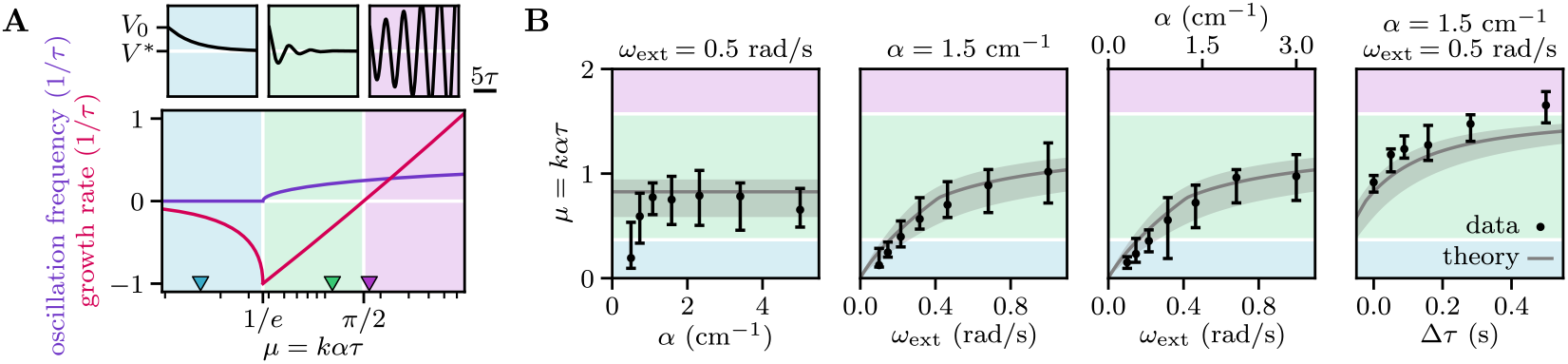
Fish adapt their responsiveness to prevent delay-induced instability. (A) Growth rate and oscillation frequency of the solutions to Eq. 1 as a function of the dimensionless gain *µ*. The three dynamical regimes are illustrated by example trajectories (top plots) for 3 values of *µ* (colored triangles). (B) Estimated values of the dimensionless gain *µ* through linear regression (black error bars, quartiles of the distributions over *N* = (33, 31, 28, 14) fish, from left to right). The predictions of the effective linear theory of Eq. 2 are shown in gray.

This simple linear model accounts for non-trivial aspects of the system dynamics. We forced speed oscillations with a sinusoidal modulation of the external flow. Using the model, we could quantitatively reproduce the observed phase lag between the fish speed oscillations and the driving signal over a wide range of driving frequencies (SI section 2.3). We also manipulated the sensorimotor delay *τ* by adding a variable delay

Δ*τ* to the visual feedback. Again, the model quantitatively captures the decrease in frequency of the self-sustained oscillation with increasing Δ*τ* (SI section 2.4). Notably, for large additional delays (*∼* 0.5 s), the oscillations become highly regular in both amplitude and frequency, as expected for a limit cycle, suggesting that the gain *µ* has crossed the bifurcation threshold.

### Adaptive responsiveness prevents delay-induced instability

The three dynamic regimes illustrated in Fig. 4A highlight the challenge that the fish need to overcome: they must operate at large enough gain *µ* = *kατ* to swiftly reach the target speed, but keep clear of the bifurcation threshold beyond which oscillations become unstable. This seems challenging considering that *µ* is proportional to the visual feedback gain *α*, a parameter that can vary rapidly over a wide range and cannot be directly inferred from visual cues.

As its value governs the system dynamics, we sought to evaluate *µ* across experimental conditions, i.e. for different values of *ω*_ext_, *α*, and of additional feedback delay Δ*τ*. Since the delay *τ* was found to be invariant, we estimated the responsiveness *k* through linear regression between 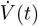 and *ω*(*t − τ*).

Assuming the responsiveness *k* to be fixed, the dimensionless gain *µ* should be proportional to both *α* and *τ*. In contrast, we found that *µ* is quasi-independent of *α* (except for target speeds approaching the upper physiological limit) and increases sublinearly with both Δ*τ* and *ω*_ext_ (Fig. 4B). These observations suggest that the responsiveness *k* is adaptive. Interestingly, across most of the range of parameters tested, the gain adaptation leads the fish to operate within the intermediate dynamic regime, which optimally balances responsiveness and stability.

### Logarithmic transformations balance responsiveness and stability

To better understand the mechanism underlying the adaptive behavior, we monitored the fish transient acceleration following a sudden change in optic flow rate. After a 10 s period during which the fish adjusted their swimming speed to match a constant optic flow rate *ω*_ext_, we suddenly disabled the feedback and displayed forward or backward flows of various rates *ω* (Fig. 5A). We found that, after a delay *τ*, the swimming speed approximately follows an exponential evolution, with an acceleration rate 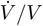 independent of both *ω*_ext_ and *α* (SI section 2.6). For |*ω*| *> ω*_th_ *≈* 0.03 rad/s, the acceleration rate was found to vary logarithmically with *ω* for both forward and backward flow, with the fish accelerating and decelerating, respectively. For |*ω*| *< ω*_th_ the fish shows a slight deceleration independent of *ω*.

**Figure 5:**
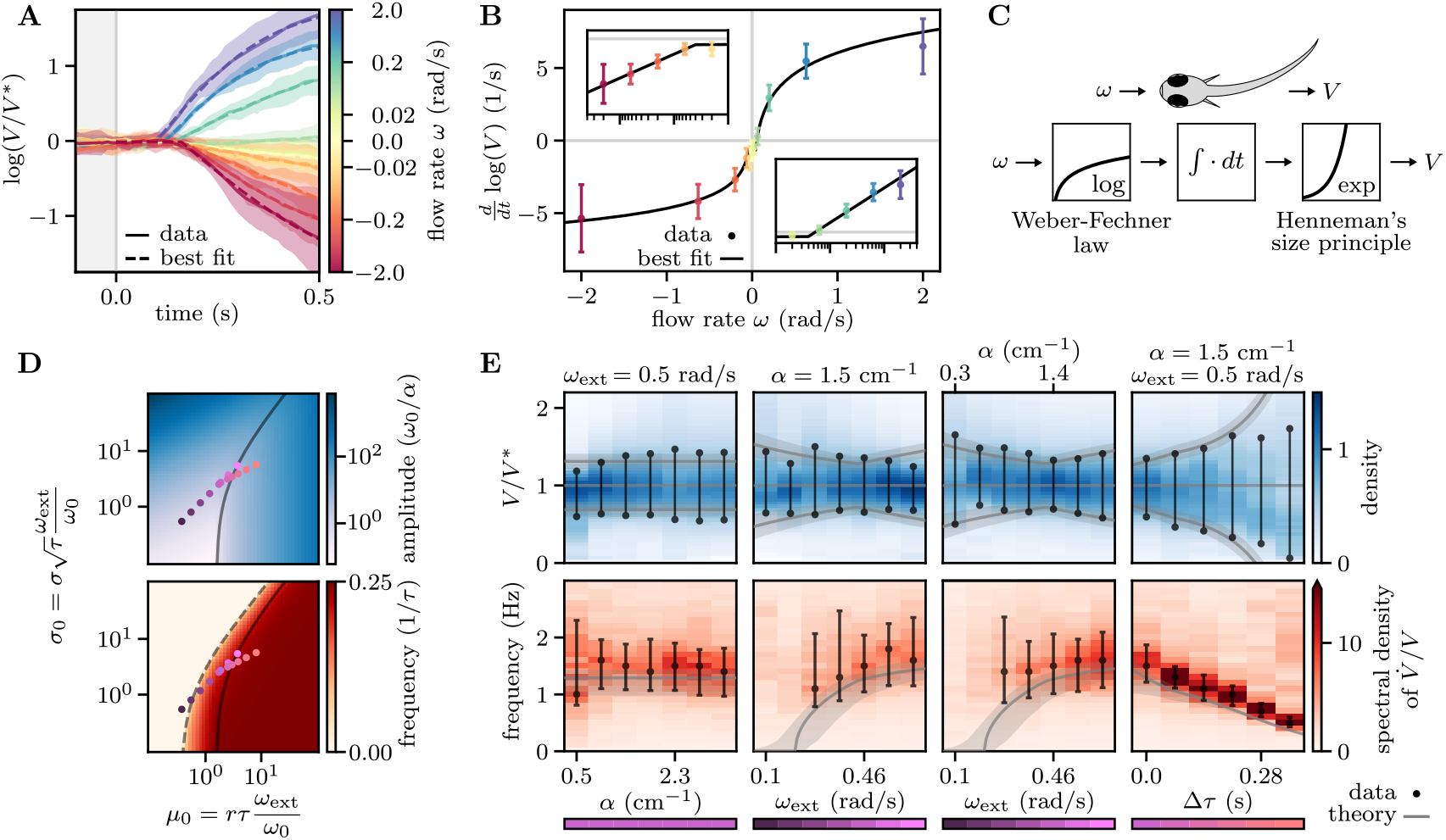
Logarithmic transformations balance responsiveness and stability. (A) Evolution of the swimming speed following a sudden perturbation, triggered at time *t* = 0, of the optic flow rate *ω* (color-coded values). Median trajectories over trials (solid lines and shaded areas, mean and standard deviation across *N* = 5 fish) evolve approximately exponentially (dashed lines, best fit) after a delay *τ*. (B) Acceleration rate (extracted from (A)) as a function of *ω*. Beyond a threshold, the dependence is logarithmic (black line, best fit). The insets show the same data on semilogarithmic scales for positive and negative *ω*. (C) Schematic showing the sequence of sensorimotor transformations. The optic flow undergoes a logarithmic compression, followed by an integration process and, finally, an exponential expansion sets the swimming speed. (D) Amplitude and frequency of the oscillations derived from Eq. 2 as a function of the dimensionless parameters *µ*_0_ and *σ*_0_. There is a sharp transition to oscillatory behavior (dashed line) and a smooth transition between noise-induced and limit cycle oscillations (solid line). (E) Top: distributions of relative swimming speeds *V/V* ^*\**^ (blue scale). The values at half-maximum (black dots) delimit the typical range of speed fluctuations. Bottom: power spectra of the acceleration rate 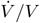 (red scale) and characteristic oscillation frequency (black error bars, for peak-to-width ratios larger than 3). Data was pooled across the fish from the experiments of Fig. 4B (*N* = (33, 31, 28, 14), from left to right). The color bars on the bottom correspond to the points in (D), leading to the theoretical predictions shown in gray.

These findings can be summarized as a logarithmic compression of the sensory input, followed by a temporal integration and an exponential expansion (Fig. 5C), and so the swimming speed can be expressed by the following stochastic nonlinear delay equation:

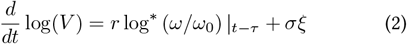

where log^*\**^ denotes the symmetrized logarithm defined as log^*\**^(*x*) = sgn(*x*) log(1 + |*x*|), a functional form that offers a reasonable approximation of the measured response function (Fig. 5B). The response parameters *r* and *ω*_0_ were inferred to be *r* = 1.6 *±* 0.3 s^*−*1^ and *ω*_0_ = 0.07 *±* 0.03 rad/s from fitting the data of Fig. 5B. A white noise source *ξ* with amplitude *σ* was also introduced to account for the additive noise observed in the dynamics of log(*V*), in agreement with the motor control literature [16]. From the variability in the stabilization to constant currents we estimated the noise amplitude at *σ* = 0.9 *±* 0.2 s^*−*1*/*2^ (Methods). Importantly, the three parameters *r, ω*_0_ and *σ* were found to be independent of the experimental conditions and weakly variable across individuals.

To better interpret Eq. 2, we can define the effective responsiveness as 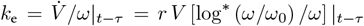. For small deviations around the target speed *V ≃ ω*_ext_*/α*, the effective gain becomes *µ*_e_ = *k*_e_ *α τ ≃ r ω*_ext_ [log^*\**^ (*ω/ω*_0_) */ω*] | _*t − τ*_. Hence, the exponential expansion results in *µ*_e_ being independent of *α* and proportional to *ω*_ext_. Furthermore, because of the logarithmic compression of the optic flow rate, *k*_e_ keeps decreasing as the swimming speed deviates from the target, preventing the oscillations from diverging.

As numerical simulations of Eq. 2 quantitatively reproduced the experimentally observed speed oscillations (SI section 2.9), we developed an effective linear theory to get an analytical understanding of the system’s behavior (Methods). In the limit of small oscillations, for which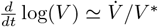, we found that the dynamics are fully determined by the dimensionless gain *µ*_0_ = *rτω*_ext_*/ω*_0_ and noise amplitude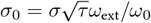.

We derived theoretical predictions for the amplitude and characteristic frequency of the fluctuations by combining the contributions of noise and nonlinearity (Fig. 5D). The deterministic dynamics (i.e. in the absence of noise) give rise to limit cycle oscillations for *µ*_0_ *> π/*2, whose amplitude corresponds to an effective responsiveness *k*_e_ close to the stability boundary *µ*_e_ = *π/*2. The presence of noise drives the system away from the deterministic attractor, leading to larger values of |*ω*| and correspondingly smaller values of *k*_e_. As in the analysis of Eq. 1, we found a sharp transition to oscillatory dynamics for *µ*_e_ = 1*/e*. However, the noise tends to smooth out the bifurcation to limit cycle oscillations at *µ*_0_ = *π/*2 (SI section 2.8).

While the value of *α* has no effect on the behavior of the solutions, increasing *ω*_ext_ or *τ* moves the system from the noise- dominated regime to the limit cycle-dominated regime with the oscillation frequency transitioning from 0 to 1*/*(4*τ*). The relative amplitude of the fluctuations depends only weakly on *ω*_ext_ but it is strongly modulated by *τ*. Plugging in our estimates of *τ, σ, r* and *ω*_0_, we obtained theoretical predictions that quantitatively match the amplitude and frequency of the measured speed fluctuations (Fig. 5E). We also found that the predicted effective responsiveness matches our naive estimates through linear regression (Fig. 4B).

### Logarithmic coding in the brain

The behavioral experiments allowed us to establish the sensorimotor operations that govern speed stabilization (Fig. 5C). We next asked whether these non-linear transformations could be reflected in the neural coding of optic flow and swimming speed. To address this, we leveraged the small size and transparency of danionella’s brain to perform brain-wide functional recordings in 2-week-old transgenic larvae expressing the GCaMP6s calcium indicators in all neurons. The reporter expression was confined to the neuronal nuclei and we used a light sheet microscope with confocal line detection in order to mitigate possible cross-talk (Methods) [17, 18].

After neuronal segmentation and signal extraction, we obtained, for each neuron, a time-trace of the relative variation in fluorescence Δ*F/F*_0_. Although we could monitor calcium activity in closed-loop settings (see movie S4), we mostly used open-loop experiments as they allowed us to disentangle visually-driven from motor-related activity. The fish were exposed to forward and backward flows of varying rates and we simultaneously recorded their tail movements. Visually-driven neurons were categorized based on their correlation (positive or negative) with forward and/or backward flow rates (Fig. 6A-B, Methods). Similarly, motor-related neurons were either positively or negatively correlated with swimming speed (Fig. 6C-D). These distinct populations displayed stereotypical spatial distributions (Fig. 6A and C, movies S5-6).

**Figure 6:**
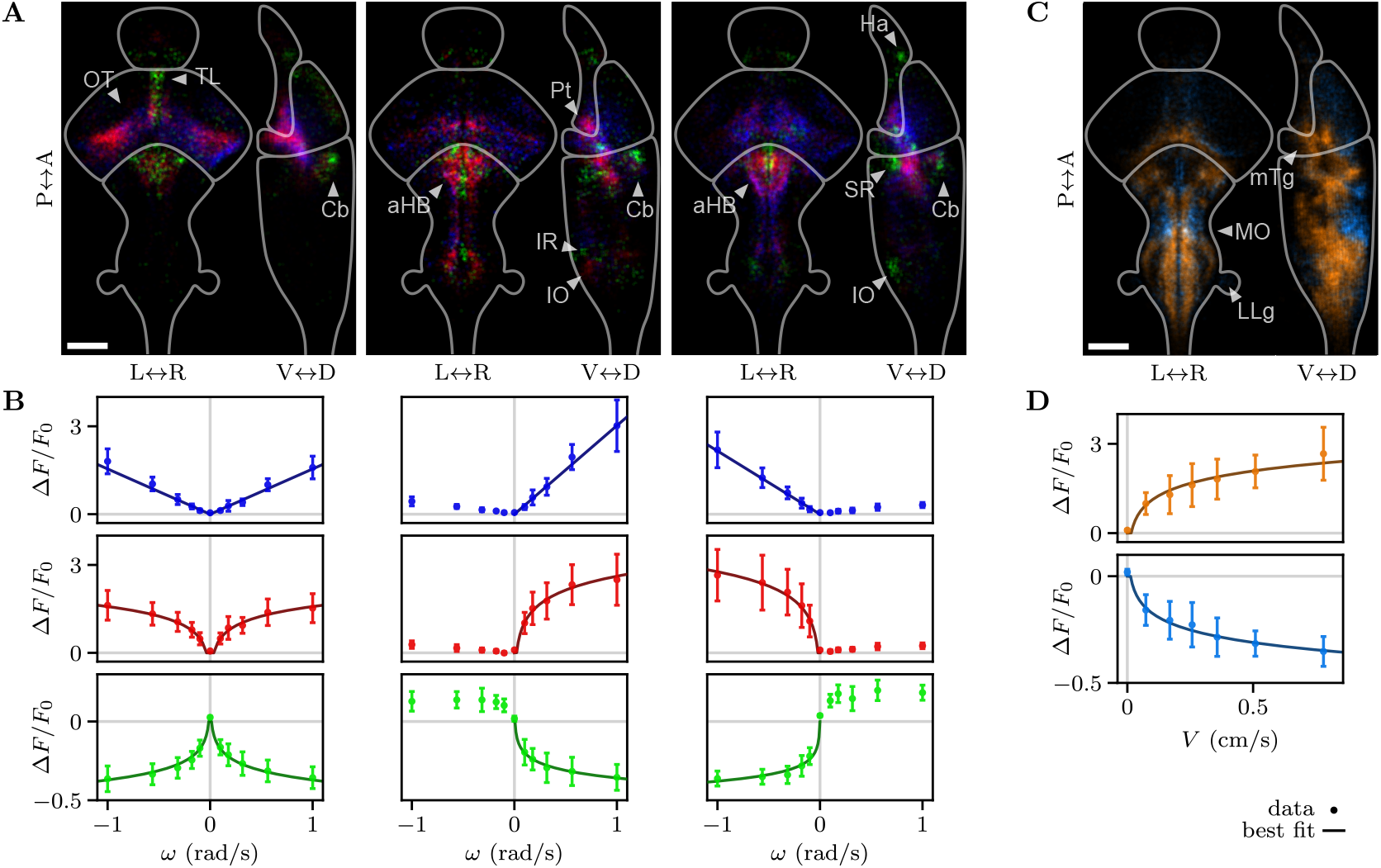
Neural activity encodes optic flow and swimming speed logarithmically. (A) Spatial distributions of neurons whose activity is correlated to optic flow rate *ω*, either to both forward and backward flow (left), only forward (middle) or only backward (right). Blue (correlated) populations show a linear response to optic flow rate, whereas red (correlated) and green (anticorrelated) are logarithmically tuned. The density distributions are averaged over *N* = 8 fish and projected in the horizontal (left) and sagittal (right) planes. They are separately normalized for each neuronal population. The outlines indicate the forebrain, midbrain and hindbrain (from top to bottom). (OT) optic tectum; (TL) torus longitudinalis; (Cb) cerebellum; (aHB) anterior hindbrain; (Pt) pretectum; (IR) inferior raphe; (IO) inferior olive; (Ha) habenula; (SR) superior raphe. Scale bar: 100 µm. (B) Tuning curves for the populations shown in (A), showing the average activity of each population as a function of *ω* (error bars, mean and standard deviation across *N* = 8 fish) and the best fits (solid lines). (C) Same as (A), but for neurons encoding the swimming speed *V*, either increasing (orange) or decreasing (light blue) logarithmically. (MO) medulla oblongata; (mTg) medial tegmentum; (LLg) lateral line ganglia. (D) Same as (B), but for the average activity of the populations shown in (C) as a function of *V*. The solid lines show the logarithmic fits.

In line with previous observations made in zebrafish [19,20], cells positively correlated with optic flow were found in the optic tectum, pretectum, and anterior hindbrain (Fig. 6A). We also identified neuronal populations anti-correlated with optic flow in the habenula, torus longitudinalis, cerebellum, superior and inferior raphe, and inferior olive. We further categorized these neurons based on the shape of their tuning curves (Fig. 6B). We observed that, as the information driving the optomotor response propagates from the visual regions to the hindbrain, the tuning undergoes a transformation from mostly linear to logarithmic coding. We checked that these nonlinear response curves could not be accounted for by a saturation of the calcium sensor (SI section 3.6).

Neuronal populations encoding the swimming speed were distributed throughout the hindbrain (Fig. 6C), in regions consistent with known pre-motor areas in zebrafish [21, 22], the medulla oblungata and the medial tegmentum. We found neurons anticorrelated with swimming speed in the habenula, optic tectum, dorsal medulla, superior and inferior raphe, and lateral line ganglia. All motor-related neurons were logarithmically tuned, either positively and negatively, to the swimming speed (Fig. 6D).

In summary, the tuning curves of the neuronal populations engaged in visuomotor processing reveal nonlinear transformations along the visuomotor pathways that are consistent with the behavioral model.

## Discussion

We experimentally probed the sensorimotor computation at play during visually-driven speed regulation in danionella, by systematically manipulating the sensory feedback. This allowed us to infer a simple mathematical model that captures the speed of animal, including the sustained oscillations induced by the sensorimotor delay, across a wide range of experimental conditions. Our analysis reveals that the nonlinear transformations that occur at both sensory and motor ends, prevent instability and enable efficient gain adaptation.

These sensorimotor operations, inferred from behavioral assays, were evident at the neural level in the tuning curves of visually-driven and motor-associated neurons. They are also biologically grounded, reflecting two well-established principles in neuroscience (Fig. 5C). The sensory compression seen in our data is an instance of the Weber-Fechner law, which states that the perceived intensity of a stimulus increases logarithmically beyond a certain detection threshold [23]. The expansion observed at the motor end corresponds to Henne- man’s size principle [24], which states that as the motor command increases, progressively larger motor units are recruited, resulting in an exponential transformation from motor command to muscle force [25].

Zebrafish has been an important model system to study visuomotor control in vertebrates. Although the capacity of this animal to adapt its response to changes in the feedback gain is now well documented, the control mechanism at play, and notably the existence of an internal model, remains debated [26–29]. The difficulty in reaching a definitive conclusion arises from the intermittent nature of zebrafish locomotion, which obscures the underlying sensorimotor computation. Given the close morphological and genetical proximity between danionella and zebrafish, and the existence of a comparable sensorimotor delay in both species [28], it seems likely that they share the same control mechanism. In the case of zebrafish however, the integrated optic flow rate is not directly reflected in the swimming speed, but we expect it to be encoded in the persistent activity of neuronal circuits that modulate the intensity, frequency and duration of swimming bouts [30].

Because it relies on fundamental neural encoding principles, we expect the control mechanism identified in danionella to be relevant across other species and sensorimotor systems in which responsiveness and stability need to be balanced. These include flight control in insects [31,32], ocular pursuit in monkeys [33], motor control in humans [34] and position stabilization in electric fish [35], all examples in which latency-induced speed oscillations have been observed. Beyond biology, the proposed mechanism could also inspire the development of robust control strategies for robots navigating unpredictable environments [36].

## Materials and Methods

### Virtual reality experiments

All experiments were performed on larvae of *Danionella cerebrum* of age *≈* 14 days post fertilization and total body length *≈* 4.5 mm (SI section 1.1). The experimental procedures were approved by the ethics committee “Le Comité d’Éthique pour l’Expérimentation Animale Charles Darwin C2EA- 005”. Imaging experiment were performed using the transgenic line *Tg(elavl3:H2B-GCaMP6s)* [11], while behavioral experiments were also performed on wild type fish. The fish were immobilized in a 2% agarose gel (SI section 1.2) and placed in a custom-made tank (SI section 1.3). A Python program was written to compute the fictive velocity from the recorded tail movements and update the projected visual pattern in real-time (SI section 1.5). We used fluid dynamics to estimate the thrust and drag acting on the fish and computed its fictive forward speed *V* and angular velocity *Ω* (SI section 1.4). We calibrated these estimates using recordings of freely swimming fish. We projected a random pattern with a characteristic lengthscale of 5 mm on the bottom of the tank at a distance of *h* = 5 mm from the fish. To restore visual feedback we translated the pattern opposite to the fish heading direction with speed *αV h* and rotated it with angular velocity *βΩ*. To simulate an external current we translated the pattern with speed *ω*_ext_*h* along a direction that rotated together with the pattern, corresponding to a fixed direction in the virtual environment (SI section 1.5). To study the response to constant currents we presented currents with different parameter values in a randomized order, with each value presented in two trials of 30 s duration.

### Impulse response estimation via reverse correlation

We presented fish with an external current of 5 minutes duration with a fluctuating flow rate of the form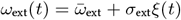, where *ξ* is a Gaussian white noise. Then the impulse response function *Ġ* was estimated by computing the cross-correlation [15] (SI section 2.2):

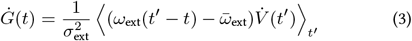

Where we averaged over the periods of time where the fish were swimming. The impulse response could be reproduced (red line in Fig. 3B) by considering the first positive peak (yellow area in Fig. 3B, Gaussian fit) to be the response kernel *K* and numerically integrating the equation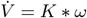.

### Analysis of linear delay equation

We nondimensionalized Eq. 1 by considering *x*(*t*^*′*^) = *V* (*t*) *− ω*_ext_*/α* and *t*^*′*^ = *t/τ*, leading to (SI section 2.1):

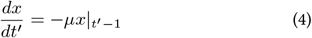

Looking for a solution in the form of an exponential with complex growth rate *λ* we obtain the following characteristic equation:

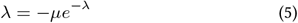

Whose solutions are given by the Lambert *W* function as *λ*_*l*_ = *W*_*l*_(*−µ*). The behavior is dominated by the largest growth rate Re(*W*_0_(*−µ*)), with the corresponding oscillation frequency Im(*W*_0_(*−µ*))*/*(2*π*).

### Responsiveness estimation via linear regression

We extracted the slope of the relationship between *ω*(*t − τ*) and 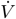 using the Theil-Sen estimator to reduce the effect of outliers [37] (SI section 2.5). We used the estimate of *τ* obtained from the impulse response and considered the data from the experiments were the fish were swimming against constant currents.

### Open-loop perturbations

We presented fish with constant currents in closed-loop and then suddenly presented a certain optic flow rate *ω* in open-loop for a duration of 0.5 s. The different values of *ω* were presented in a randomized order with each value presented at least 10 times for each one of several baseline closed-loop conditions (*ω*_ext_, *α*) (SI section 2.6). The evolution of the speed following the perturbation was fit with a delayed exponential evolution:

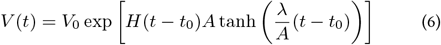

Where *H* is the Heaviside step function and we included a saturation in the form of a hyperbolic tangent. The dependence of the acceleration rate *λ* on the flow rate was fit with a logarithm:

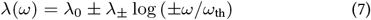

Respectively for positive and negative values of *ω*.

### Noise amplitude estimation

We estimated the noise amplitude by computing the standard deviation of the increments of the speed on a logarithmic scale (SI section 2.7):

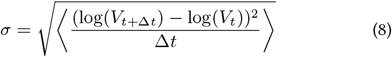

For the estimate we considered the data from the experiments were the fish were swimming against constant currents.

### Analysis of stochastic nonlinear delay equation

We nondimensionalized Eq. 2 in the limit of small oscillations by considering *x*(*t*^*′*^) = (*ω*_ext_ *− αV* (*t*))*/ω*_0_ and *t*^*′*^ = *t/τ*, leading to (SI section 2.8):

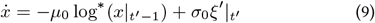

Where *ξ*^*′*^ denotes delta-correlated noise in the dimensionless time *t*^*′*^. For a certain value of 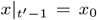 the system evolves locally with a rate of change *−µ*_0_ log^*\**^(*x*_0_) which would equivalently arise from a linear system with a gain *µ*_e_ = *µ*_0_ log^*\**^(*x*_0_)*/x*_0_. We obtained an estimate of the amplitude of limit cycle oscillations *A*_lc_ by considering the value of *x*_0_ for which the effective gain becomes *π/*2:

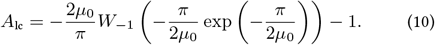

The noise term induces fluctuations with an amplitude *A*_n_ that can be derived from the following implicit relation:

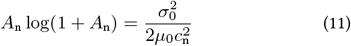

Which we obtained using the fact that a Ornstein-Uhlenbeck process with gain *µ*_e_ and noise amplitude *σ*_0_ leads to a fluctuations with standard deviation 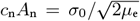 [38]. By combining the limit cycle and the noise-induced amplitudes we obtained an estimate of the total amplitude of the fluctuations and thus of the effective gain *µ*_e_(*µ*_0_, *σ*_0_). The predicted oscillation frequency is then given by Im(*W*_0_(*−µ*_e_))*/*(2*π*).

### Brain imaging experiments

We used a custom-made light sheet microscope with confocal slit detection to excite and record the fluorescence across the brain of the fish (SI section 3.1). We recorded stacks of 25 different layers at a rate of *≈* 2.3 brain volumes per second for a laser power of *≈* 150 µW. For the open-loop experiments we presented constant currents with different values of flow rate *ω* in a randomized order, with each value presented in 3 trials of 20 s duration (SI section 3.2).

### Processing of brain images

We corrected an eventual horizontal shift in the images by computing the cross-correlation between different images (SI section 3.3). Then we computed the average covariance between the intensity of neighboring pixels over the recording. Applying a blob detection algorithm to the resulting images gave the positions of all the neurons showing a modulation in activity during the recording. Then we computed the relative fluorescence change Δ*F/F*_0_ for each neuron taking as baseline *F*_0_ the median fluorescence in the absence of optic flow and tail movements.

### Registration to a reference brain

To compare the results for different fish we acquired a stack of brain images with high vertical resolution and long exposure for each fish. Then we numerically estimated the optimal affine transform between different stacks by maximizing their cross-correlation through gradient descent (SI section 3.4). The reference stack was obtained by averaging the aligned stacks for all fish and then mapping the left-right symmetry plane to the middle plane of the stack.

### Correlation analysis

We identified the neurons whose activity is significantly correlated or anticorrelated with four different signal (SI section 3.5): flow rate |*ω*|, forward flow rate *−ωH*(*−ω*), backward flow rate *ωH*(*ω*), and swimming speed *V*. We computed the Pearson correlation between these signal and the relative fluorescence change Δ*F/F*_0_ for each neuron. To determine significantly high correlation values we built null distributions by computing the correlation with time-shifted versions of the signals with an autocorrelation smaller than 0.25 in magnitude. To account for the problem of multiple comparison for hypothesis testing we used the BenjaminiHochberg procedure [39], controlling the false discovery rate with a significance threshold of 10^*−*5^. Neurons present in more than one out of the 8 identified populations were assigned to the population for which the correlation was largest in magnitude. To separate the neurons correlated to optic flow into subpolations showing linear or logarithmic coding, we computed the correlation values with the logarithm of the corresponding signal and compared them with the previously computed values. We fitted the tuning curves either with a linear response *Ax* or a logerithmic one *A* log(*x/x*_0_)*H*(*x− x*_0_) as a function of the corresponding signal *x*. To visualize the spatial distributions of the neuronal populations we used kernel density estimation with a uniform spherical kernel of radius 5 µm. We assigned a color to each pixel by computing the weigthed sum of the hues corresponding to each population with the respective value of the density. We separately normalized each density to to result in a maximum pixel intensity of 1, and the weighted sum was renormalized when any color channel exceeded 1. We used the Max Planck Zebrafish Brain (mapzebrain) atlas [40] to identify the brain regions associated with the neuronal populations.

## Materials and data availability

Data and code have been deposited in a public repository on Zenodo [41].

## Acknowledgments

We thank the members of the IBPS aquatic animal facility for their help in taking care of the fish. We thank Benjamin Judkewitz and Filippo Del Bene for sharing their fish with us and Julie Lafaye for helping set up the facility. We thank Pavan Kaushik, Verity Cook, and the members of the Laboratoire Jean Perrin and the SmartNets ETN for helpful feedback and discussions. This work was supported by the European Union’s Horizon 2020 research and innovation programme under the Marie Skłodowska-Curie grant agreement No. 860949 and by the Agence Nationale de la Recherche under the Locomat project ANR-21-CE16-0037.

## 1 Virtual reality system

### 1.1 Animal care

All the experimental procedures involving fish (*Danionella cerebrum*) were approved by the ethics committee “Le Comité d’Éthique pour l’Expérimentation Animale Charles Darwin C2EA-005” (APAFIS authorization number: 52420-2024121210531432 v7). Adult fish are kept in a system with circulating water in groups of *≈* 50 individuals in tanks of 30 L volume. The water temperature is kept at *≈* 27^*°*^C, the pH at *≈* 7.3 and the conductivity at *≈* 330 µS/cm. The light in the room follows a day/night cycle of 14/10 hours. The tanks are monitored daily to collect newly laid egg clutches. These are moved to Petri dishes with E3 medium and kept at 28^*°*^C in an incubator. After hatching, the larvae are moved to beakers of 1 L and fed twice daily with rotifers until the age of two weeks (*±*1 day), at which point they are either used for experiments or transferred into the system in tanks of 3 L. We used a wild type line and a transgenic line [1] *Tg(elavl3:H2B-GCaMP6s)* which expresses the calcium indicator GCaMP6s [2] bound to the histone H2B in the majority of neurons, because of the elavl3 promoter. The calcium indicator is localized to the nuclei of the neurons because of the interactions between the histone and the DNA. Fish from both lines were used for the behavioral experiments, whereas only transgenic fish were used in the imaging experiments.

### 1.2 Head immobilization protocol

A fish aged 14 days post fertilization is anesthetized by placing it in a solution of tricaine (*≈* 150 mg/L) buffered to pH 7 using sodium bicarbonate [3]. After about 3 minutes, it is moved in a small drop of solution on a clean PMMA surface using a transfer pipette. A drop (*≈* 1 mL) of low-melting point agarose solution (2%) at *∼* 40^*°*^C is poured on top of the fish. The fish is immediately pulled head first into a glass capillary using a piston (Drummond Wiretrol II Calibrated Micropipets, 50–100 µL), where it is left for about one minute to let the agarose solidify. At this point the capillary is inserted into a small custom-made tank, where the fish in his agarose rod can be extruded into E3 medium, allowing the oxygen to diffuse through the gel and reach the fish. The agarose around the tail of the fish is removed with the help of a scalpel. The cut is done midway between the pectoral fins and the cloaca. In order to do so, the fish is pulled halfway inside the capillary where the cut is to be made, so that the glass opposes resistance and the scalpel can cut through the agarose without deforming the rod.

### 1.3 Design of the experimental setups

We used two distinct virtual reality setups to collect the data. They function in the same way, but one is only used to record the behavior, whereas the other has been modified to enable simultaneous functional imaging. Both setups include a computer connected to a camera (FLIR Chameleon3 USB3 CM3-U3-13Y3M, with macro lens Navitar Zoom 7000) recording images of the tail and a projector (Optoma UHD35x) displaying a visual pattern on the bottom of a water tank.

The first behavioral setup consists of a tank (Fig. S1A) whose walls are laser cut rectangular blocks of PMMA, glued together with a silicon-based glue (Bostik FIXPRO MSP 111). The sides are black whereas the bottom is transparent. However, the opaque white sticker protecting the material surface is left on the upper side of the tank bottom to act as a screen for the projector. We chose to place the screen inside the tank to avoid the optical distortions and limited field of view caused by refraction at the air water interface if the screen had been placed on the outer surface of the tank [4]. The projector is placed beneath the tank and the camera is positioned above it. We also placed an IR led (OSRAM LZ4-40R608) beneath the tank to illuminate the fish and an IR filter (Thorlabs FGL780S) in front of the camera objective to block the light coming from the projector. A hole, drilled through the side of the tank at a height of 5 mm above the bottom, allows us to insert the fish in its glass capillary inside the tank. The bottom of the tank is a 5 *×* 5 cm square. With this configuration, the screen covers most of the lower visual field of the fish, up to a visual angle of *≈* 80^*°*^. As the tank is elevated on a platform, we position a mirror at 45^*°*^ underneath it to redirect the light coming from a projector at a distance of *≈* 30 cm. We set the projector contrast to *−*40, corresponding to an irradiance of *≈* 0.3 W/m^2^ measured right after the screen for a uniform white image.

The tank used for simultaneous functional imaging (Fig. S1B) is adapted from the behavioral one to accomodate a lightsheet microscope. It is slightly smaller (side length of 4.2 cm) to allow space for positioning the illumination objective on one side. The lateral walls are thin glass microscope slides to allow the laser light to propagate through the sample, and the IR light of a LED (OSRAM SFH 4545) to illuminate the fish from the opposite side. Because of the presence of the fluorescence detection objective above the tank, the behavioral camera is positioned below the tank. A hot mirror (Thorlabs M254H45) is used to redirect the IR light to the behavioral camera while directing the visible light from the projector to the screen. A small transparent window at the bottom of the tank, created by removing a 5 *×* 5 mm of the white sticker below the tail, enables monitoring the tail movement from below.

We realized that the upper half of the visual field plays an important role in the optic flow perception. In the behavioral setup the fish sees the pattern reflected on the water surface thanks to the total internal reflection, whereas in the light sheet setup this effect is perturbed by the presence of the water-dipping detection objective positioned right above the fish. We thus place a small mirror between the fish and the objective to restore the effect of the water surface reflection and increase the fraction of visual field that is stimulated. The mirror is custom-made using a thin glass slide (thickness *≈* 100 µm), which we paint with a spray to make it reflective (MTN PRO Glass to Mirror Converter) and coat in black (MTN 94 Black). To enable fluorescence brain imaging from above, we create a transparent window (diameter *≈* 2 mm) by covering the slide with a round sticker before spraying it. The mirror is placed *≈* 1 mm above the fish with a support made of several pieces of transparent PMMA of various thicknesses. We include a thin holder (*≈* 250 µm) with a small hole (diameter *≈* 1 mm) to hold the agarose rod, preventing motion when the fish swims and ensuring that the neurons maintain a stable position in the brain images. The mirror support is extended above the tail and painted in black to create a uniform background for the tail. For brain imaging, we use a detection objective with a large working distance (*≈* 5.5 mm), which leaves enough space and minimizes the effect of spherical aberrations due to the presence of the glass slide.

The behavioral experiments were done on both setups and we did not notice any difference in the observed behavior.

### 1.4 Fictive velocity estimation

The problem of estimating the intended movements of head-restrained fish has been addressed in various ways by scientists who developed virtual reality setups for larval zebrafish [5–9]. Here, we decided to take a slightly different approach and tried to estimate the speed of the fish using a fluid dynamics model. In order to propel itself, danionella oscillates its tail, pushing water backwards and therefore accelerating as a result of the reactive forces exerted by the water. To estimate the thrust generated by the fish, we use a model developed by Lighthill [10]. The author considers the rate of change of momentum in a volume of water surrounding the fish and derive an expression for the thrust. The result, averaged over an oscillation period, only depends on the velocity of the tip of the tail 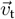 and on its orientation. As shown schematically in Fig. S2A, we take 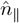 to be the unit vector tangent to the tail midline at the tip, and 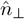 the one perpendicular to it. Denoting the components of the tail velocity parallel and perpendicular to the tail direction as 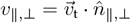, the reaction force is given by:

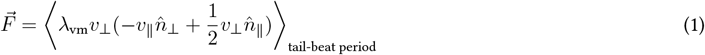

where the angular brackets indicate an average over an oscillation period and *λ*_vm_ is the virtual mass per unit length at the tip. The latter corresponds to the mass of water displaced by the tail movements and can be approximated as *λ*_vm_ = *ρπh*^2^*/*4, where *ρ* is the water density and *h* the height of the tail. The forward thrust is then given by the component of 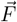 in the fish heading direction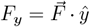. If we assume that the force is acting at the tip of the tail then the torque about the fish’s head is:

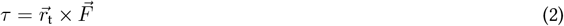

where 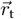 is the position of the tip of the tail with respect to the fish’s head.

### Forward speed

To compute the speed of the fish *V*, we write down Newton’s equation:

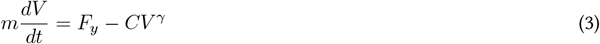

The first term on the right-hand side represents the thrust exerted by the fish to accelerate forward whereas the second term is the drag force that slows it down. There can be various contributions to the drag that are more or less relevant depending on the fluid dynamic regime. Two-week old fish, such as the ones studied here, are in an intermediate fluid dynamic regime: the Reynolds number can be estimated at *Re ≈* 25 by considering a characteristic fish speed *V*_c_ *≈* 0.5 cm/s and length *L*_c_ *≈* 0.5 cm. In this regime, the scaling exponent of the drag force is expected to be *γ* = 3*/*2 [11]. Then, for a fixed thrust the fish reaches a steady state speed *V* = (*F*_*y*_*/C*)^2*/*3^. We used this expression as an estimate of the fish speed, this last step is equivalent to ignoring the effect of inertia. We do so in order to close the feedback loop as fast as possible, as there is an incompressible delay induced by the velocity estimation and subsequent presentation of the visual feedback, which plays a similar role to inertia. Putting everything together we estimated the speed of the fish up to a proportionality constant *A* as:

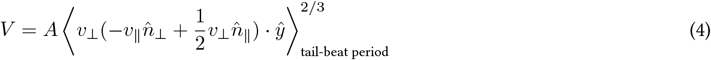

In order to test the validity of this estimate, we recorded freely swimming fish in a quasi-two-dimensional tank. To obtain a high resolution of the tail movement, we used a camera mounted on a motor to track the position of the fish in real-time. The specifics of this setup will be described elsewhere. We processed the images to extract the fish position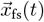 and orientation *θ*_fs_(*t*). We computed the time derivative of the position signal to estimate the fish speed *V*_fs_(*t*). Moreover, we segmented the tail with a piecewise linear curve and computed the estimate *V* from Eq. 4. We found that it correlates well with the actual speed of the fish and we extracted an estimate of the proportionality constant *A* by minimizing the mean squared error:

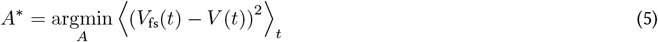

Fig. S2D shows both *V*_fs_(*t*) and *V* (*t*) inferred from the tail movement, for one example fish. illustrating the accuracy of this estimation. In Fig. S2B we plotted the distribution of values of *V*_fs_(*t*) and *V* (*t*) pooled over *N* = 3 fish. Although most of the measurements are close to the identity line, one can see a secondary peak that deviates from the identity. This bimodality might be due to the fact that the fish do not always maintain an horizontal posture. They sometime display a tilted (nose-down) position, dragging their head against the bottom of the tank. Nonetheless, we calculated the Pearson correlation coefficient between the two signals and found a reasonably high value of *≈* 0.6. We also found that the value of *A*^*\**^ is consistent across the different fish with an average of *A*^*\**^ *≈* 0.2 (cm/s)^1*/*3^. We measured the time shift between *V* (*t*) and *V*_fs_(*t*) by computing the cross-correlation between the two signals, we found that *V*_fs_(*t*) follows *V* (*t*) with a delay Δ*t*_*V*_ = 22 *±* 5 ms, giving an estimate of the inertial timescale that we neglected in the calculation.

### Angular velocity

To estimate the fish angular velocity *Ω*, we write down the rotational form of Newton’s equation:

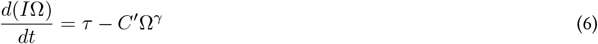

where *I* is the moment of inertia of the fish, and the second term on the right-hand side is a drag term analogous to that of Eq. 3. One additional complication arises from the fact that the fish’s moment of inertia changes as it turns [12]. Moreover, as the fish is not a rigid object - it deforms as it turns - the form of the drag force chosen in Eq. 6 is also an oversimplification. Still we can derive an estimate for the angular velocity analogously to how we did for the speed. We ignore inertia and consider the regime of viscous drag *γ* = 1 to get *Ω* = *τ/C*^*′*^. Thus, subtituting the expression for the torque, we get the angular velocity up to a proportionality constant *B*:

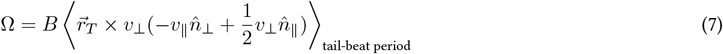

Using the freely swimming data we computed both *Ω* from Eq. 7 and the actual angular speed of the fish *ω*_fs_. By examining the cross-correlation between the two signals, we found that *ω*_fs_ actually precedes *Ω* with an average time shift of Δ*t*_*Ω*_ = *−*15 *±* 3 ms. This suggests that the dynamics of the angular velocity is fast and it justifies *a posteriori* our quasi-steady state approximation and the choice of viscous drag *γ* = 1. We observed that the fish perform discrete turning events of *≈* 200 ms duration. We are not interested in capturing the fine scale details of the angular velocity but rather the net reorientation angles associated with these turns, therefore we considered the orientation difference between *t* and *t* + *δt*:

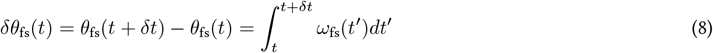

This corresponds to the total reorientation angle in the following time interval of duration *δt*. We compared the measured orientation difference *δθ*_fs_(*t*) with the integral of *Ω*(*t*) and found that they are well correlated. As for the forward speed, we computed an estimate of the proportionality constant *B* by minimizing the mean squared error between the two signals:

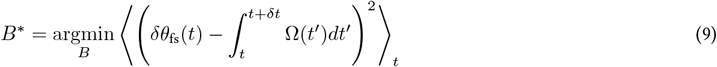

In the computation of the reorientation angle, we choose *δt* = 1 s, a value that is larger than the typical duration of a single turning event but also smaller than the time interval between successive turns so that the effect of multiple turns does not overlap in the computed signal. In Fig. S2E we plotted 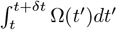 together with *δθ*_fs_(*t*) for one example fish. For most turning events, the net reorientation angle is well captured by our estimation computed from the tail movements alone. In Fig. S2C, we plotted the distribution of these quantities pooled over *N* = 3 fish. We found that most of the measurements are closed to the identity line, confirming the quality of the estimation method. The Pearson correlation coefficient between the two signals was found to be *≈* 0.8. As part of this correlation comes from the sign of the reorientation angles, we further assessed how well the magnitude of the reorientations is captured. We thus computed the correlation coefficient by considering the absolute values of the signals and still found a high value of *≈* 0.7. The value of *B*^*\**^ is also consistent across different fish with an average of *B*^*\**^ *≈* 200 deg·s/cm^3^.

### 1.5 Virtual reality implementation

A Python program was written to capture images from the camera (using the FLIR PySpin library), and display visual patterns with the projector (using the PsychoPy library [13]). Tail images with a pixel size of *≈* 30 µm were acquired at a frame rate of 200 Hz. For each image, we segmented the tail with a piecewise linear curve by starting from a point close to the head and iteratively choosing the next point in the direction of the center of mass (weighing the pixel intensities) evaluated in a circular region centered around the expected position of the next point [14]. Then we evaluated the fictive speed *V* and angular velocity *Ω* from Eqs. 4 and 7, using multiple successive images to compute the velocity of the tail tip. We computed the time-average over all frames up to 60 ms in the past, corresponding to approximately one tail-beat period. Performing this rolling average introduced a delay of 30 ms in the visual feedback.

We used the projector to display a high-contrast random pattern and updated its position and orientation with a refresh rate of 120 Hz. We used a Turing-like pattern created by considering a Fourier spectrum with a single peak at a characteristic length scale *λ* and a Gaussian random phase. We computed the inverse Fourier transform to get the image and binarized it to maximize the contrast. The typical diameter of black and white regions in the pattern is then given by *λ/*2. We chose *λ ≈* 5 mm resulting in large enough features to be easily distinguishable from a distance of *h ≈* 5 mm.

When a fish swims forward at a certain speed, the surrounding environment appears to move in the opposite direction with that same speed. Thus we translated the pattern opposite to the fish’s heading direction 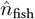 with velocity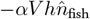, with *α* being the translational feedback gain. This translation corresponds to an optic flow rate *αV*. Similarly, if the fish turns in one direction then the environment appears to rotate around it in the opposite direction. Therefore the pattern is rotated about the fish head with angular velocity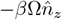, where *β* is the rotational feedback gain and 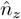 is the unit vector pointing upwards from the fish head.

The finite time needed to process the images and update the projected pattern results in an additional feedback delay. We measured the latency of the system using an additional camera recording at a 240 Hz to capture the reaction time of the visual pattern in response to an external light stimulus to the camera and found a delay of 30 *±* 4 ms. Adding this to the delay introduced by the rolling average we get a total feedback delay *τ*_f_ = 60 *±* 4 ms.

In addition to the visual feedback, we displaced the pattern independently of the fish’s actions to simulate the presence of an external current dragging the fish in a given direction. Translating the pattern with velocity 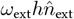 simulates a fish movement in the opposite direction with optic flow rate *ω*_ext_. The direction of the current 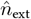 is not fixed relative the fish heading, but rather to the allocentric reference frame defined by the virtual environment. This way the fish can actually change its orientation in the virtual space and align to the external current. The update of the visual pattern in a small time interval Δ*t* is thus a rotation about the fish head of an angle *−βΩ*Δ*t* and a translation of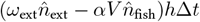.

### 1.6 Behavioral protocol

The experimental room temperature was kept at 24 *±* 1^*°*^C. The room was kept in the dark, with the only source of visible light for the fish being the projector (and the blue laser in the imaging experiments). After the head-immobilization procedure, fish were left to habituate for at least 15 minutes before proceeding with the experiments. We noted that, after this habituation period, fish rarely swam unless stimulated. We first presented a forward visual flow in open-loop conditions (*α* = 0, *β* = 0) to check whether the fish were responsive to the visual stimulus and exhibited the optomotor response. We then presented visual flows in closed-loop conditions with different values of flow rate *ω*_ext_ and feedback gain *α* to check if the fish adapted their swimming speed to stabilize the external currents. Some of the tested fish would not swim continuously, but only performed struggling bouts or swimmed at a very large speed for short periods of time. We selected for the ensuing experiments only fish that were able to modulate their speed in response to different flows. These performing animals corresponded to *≈* 62% of the tested fish. As the natural rotational gain *β* = 1 could lead to oscillations and sometimes instability of the fish orientation we decided to set *β* = 0.25 in all of the experiments, except when noted otherwise. We think that this instability is due to the fact that in freely swimming conditions the orientation changes fast in response to tail movements and the feedback delay introduced by the virtual reality can make the orientation control unstable. To test the response to different conditions (different values of *α* and *ω*_ext_) we presented currents of 30 s duration alternated with pauses of 30 s during which *ω*_ext_ = 0. At the beginning of each trial the current is aligned with the fish heading direction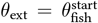. Different conditions were presented in a randomized fashion, with each condition appearing in two distinct trials.

### 1.7 Example responses in different conditions

Fig. S3 shows some examples of the fish response to external currents under various conditions. Fig. S3A shows how, in the absence of feedback, a fish typically swim at high speed and stops shortly after, whereas for an appropriate value of *α*, it swims continuously and adapts its speed to match that of the current. Fig. S3B shows the response of a fish for a target speed slightly larger than the maximum physiologically attainable value, for which it swims for a few seconds below the target before struggling and giving up. When the target speed is smaller than the minimum accessible value, the fish swims intermittently. Fig. S3C shows example trials with different external flow rates, the fish adapts its swimming speed to match that of the current on average and thus stabilize the optic flow. Fig. S3D shows example trials with different feedback gains. The fish adapts its swimming speed to stabilize the optic flow, even though the external flow rate stays the same. Fig. S3E shows example trials with currents oriented in different directions. The fish turns to reorient itself in the virtual environment and then swims against the current to stabilize its position.

### 1.8 Oscillations of tail deflection and swimming speed

In Fig. S4 we illustrate the tail oscillations that underlie swimming. We define the tail deflection as the angle *θ*_t_ between the body axis and the line connecting the base and tip of the tail (Fig. S4C). When a fish stabilizes the visual flow, its tail oscillates with an approximately constant tail-beat frequency and an amplitude that is modulated in time (Fig. S4A). Turning events to align to currents in different directions correspond to biased tail deflections to either the left or right side (Fig. S4B). We computed the Fourier spectrum of the rate of change of the tail deflection *θ*_t_ and found that it has a single peak at a frequency of 16 *±* 2 Hz (Fig. S4D). To show the different timescale of the speed oscillations that we observe during stabilization we also computed the Fourier spectrum of the relative acceleration 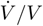 and found that it has a peak at a frequency of 1.5 *±* 0.7 Hz (Fig. S4E).

### 1.9 Swimming speed distributions

To give an idea of the scale of the speed fluctuations in different conditions we plotted the distributions of swimming speed *V* for the experiments with constant currents. We pooled together the data from all the fish for each condition, only considering the times where the fish are swimming. We see that the distributions show a single peak whose position varies depending on the condition, reflecting the fact that fish adapt their speed to the target value *V* ^*\**^ (Fig. S5). Moreover we noticed that the fluctuation amplitude, corresponding to the width of the distributions, seems to scale with the target speed. When plotting the distributions of the normalized speed *V/V* ^*\**^ we find that they approximately collapse onto the same curve, both for varying values of *α* and *ω*_ext_ (Fig. S5A,B). We find a deviation from this collapse for both small and large values of the target speed, corresponding to values close to the boundaries of the physiologically accessible range. This suggests that the amplitude of the speed fluctuations is proportional to the speed itself, or equivalently that the fluctuations of the logarithm log(*V*) have a constant amplitude. When varying both *α* and *ω*_ext_ with a constant target *V* ^*\**^ we find that the distribution becomes skewed to larger values of *V* only for a very small value of *ω*_ext_ (Fig. S5C). This effect is due to the fact that fish start swimming at a relatively high speed and then only slowly adapt to the target because of the small optic flow rate. When increasing the feedback delay, the amplitude of the fluctuation increases as shown in Fig. S5D.

## 2 Stabilization with delays

### 2.1 Linear delay equation analysis

Here we study the solutions to the linear delay equation:

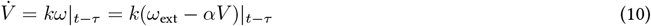

We consider the case of a slowly varying external flow rate, for which *ω*_ext_ | _*t − τ*_ *≃ ω*_ext_| _*t*_. Then, we can nondimensionalize the equation by considering the change of variables:

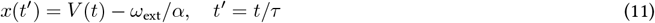

This operation shifts the dependent variable - moving the fixed point of the equation from *V* = *ω*_ext_*/α* to *x* = 0 - and rescales time so that it is measured in units of the delay *τ*. We get the equation:

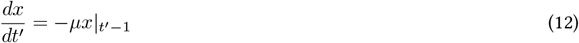

We are left with a single dimensionless parameter, *µ* = *kατ*, a dimensionless gain that combines the effect of the delay with the total sensorimotor loop gain. Delay differential equations are notoriously difficult to solve, but this one being linear, it lends itself to an analytical approach. We seek a solution in the form of a complex exponential 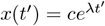 [15]. Substituting it into Eq. 12 we get:

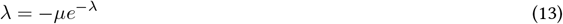

This characteristic equation is trascendental and has an infinite numbers of solutions that can be expressed in terms of the Lambert *W* function [16] as *λ*_*l*_ = *W*_*l*_(*−µ*), where *l* is an integer denoting the branch of this multivalued function. Then we can express a solution of Eq. 12 as the linear combination:

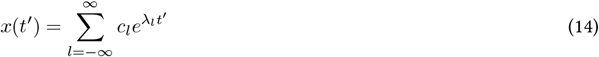

where the coefficients *c*_*l*_ are specified by the initial condition *x*(*t*^*′*^) = *x*_0_(*t*^*′*^) for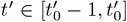]. After an initial transient the behavior of the solution is dominated by the term with the largest growth rate Re(*λ*_*l*_). This is given by *l* = 0 for *µ <* 1*/e* and both *l* = 0 and *l* = *−*1 for *µ >* 1*/e* as the two branches become complex conjugates and one gets oscillatory behavior. Then, for the growth rate we have:

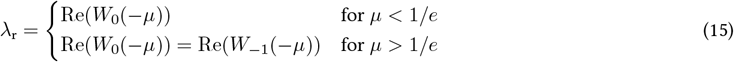

while the angular frequency is given by the imaginary part of *λ*_*l*_:

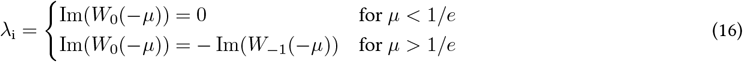

To study the stability of the solutions we have to look at the sign of the growth rate. For *µ > π/*2, *λ*_r_ *>* 0, leading to an unstable system with diverging solutions [17]. We see that it is possible to observe oscillations even if the system is one-dimensional, as the presence of the delay makes the phase space infinite-dimensional, which is also reflected in the need of a functional initial condition. As the behavior of the fixed point *x* = 0 is determined by at most two solutions of the characteristic equation, we can describe it with the nomenclature of stability theory for two-dimensional dynamical systems [18]. Then, the fixed point is a stable node (*λ*_r_ *<* 0, *λ*_i_ = 0) for *µ <* 1*/e*, a stable spiral (*λ*_r_ *<* 0, *λ*_i_ *>* 0) for 1*/e < µ < π/*2 and an unstable spiral (*λ*_r_ *>* 0, *λ*_i_ *>* 0) for *µ > π/*2.

### 2.2 Impulse response estimation via reverse correlation

Eq. 10 is a delay equation where the acceleration only depends on the optic flow rate at a single time point in the past. However, we expect the dependence to be spread out in time, partly because the optic flow rate cannot be measured instantaneously. More generally, we can expect the acceleration to depend on all previous values of the flow rate, weighted with some kernel *K*(*t*):

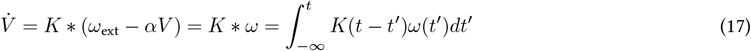

This remains a linear approximation, as there might be higher-order terms that are ignored in this expression. The canonical way to measure the kernel in a system of this sort is to inject white noise as the input (*ω*), measure the output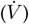, and then look at how the fluctuations of the two signals relate to each other to get an estimate of the kernel (*K*). This technique is called reverse correlation and consists in evaluating the cross-correlation between the input and the output [19]. In our system, things are a bit more complicated as we do not have complete control over the input *ω*. Indeed, this input is in part driven by the fish behavior *V* through visual feedback. We could imagine turning off the feedback, but in that case the fish would stop swimming. We can still consider a white noise perturbation of the external flow speed *ω*_ext_ and see how the fish reacts to the speed fluctuations and tries to compensate for them. Because of the linearity of Eq. 17 we can write its solution for a generic external input *ω*_ext_(*t*) in terms of the impulse response function (Green’s function). The impulse response function *G*(*t*) is the solution of Eq. 17 for *ω*_ext_(*t*) = *δ*(*t*). It allows us to write the generic solution of Eq. 17 as a convolution with the external drive: *V* = *G * ω*_ext_. Taking the time derivative we obtain an equation for the impulse response Ġ of the acceleration:

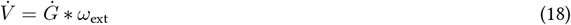

Comparing this equation with Eq. 17, we see that we can now extract the impulse response function Ġ by computing the cross-correlation between *ω*_ext_ and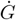. We performed experiments with an external current with a fluctuating flow rate of the form 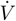 where *ξ* is a Gaussian white noise such that ⟨*ξ*⟩ = 0 and *ξ*(*t*_1_)*ξ*(*t*_2_)⟩ = *δ*(*t*_1_ *− t*_2_). Then, an estimate of *Ġ* can be obtained by evaluating:

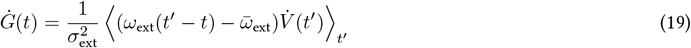

We implemented such an experimental paradigm by drawing the external flow rate from a Gaussian distribution at each update of the visual stimulus. To implement the white noise stimulus, the distribution was chosen to have a mean 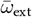 and standard deviation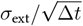, where Δ*t ≈* 8 ms is the time increment between two successive projector updates (Fig. S7A). We presented fish with such a current for a duration of 5 minutes, then we computed the cross-correlation (Eq. 19) considering only the periods during which the fish were swimming continuously. We performed this experiment for *N* = 11 fish with parameters: 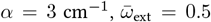 and *σ*_ext_ = 0.04 rad s^*−*1*/*2^. We found that the impulse response is comparable across different fish. It is close to zero at short times and then exhibits a sharp positive peak at a finite delay of *≈* 150 ms. This reflects an acceleration following a positive perturbation of the external current. The peak is followed by a wider and shallower negative part, reflecting a deceleration that brings the system back to the target speed. This form of the impulse response function is compatible with the linear delay model (Eq. 10), which would correspond to a kernel of the form *K*(*t*) = *kδ*(*t − τ*). The negative part of the impulse response is also expected from the model as the perturbation drives the system away from the target speed, thus changing the visual feedback and leading to a corrective motor update that restores the steady state. In other words the first peak is due to the response to the external current, reflecting the kernel *K*, whereas the following modulation reflects the internal system dynamics. This separation is possible because the response is delayed, otherwise the effects of the external current and of the internal dynamics would be mixed in the impulse response.

We fit the positive peak with a Gaussian and extracted an estimate of the delay *τ* = 150 *±* 20 ms. In principle, we could also get an estimate of the responsiveness *k* by evaluating the area under the peak, for a linear system we expect this value to remain constant independently of the scale of the fluctuations *σ*_ext_. However, we found that the value of *k* decreases with *σ*_ext_ (Fig. S7B), suggesting that the response of the system is nonlinear.

We also estimated the impulse response for different values of feedback gain *α*, average external flow rate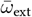, and additional feedback delay Δ*τ* (Fig. S7C). We found that the functional form remains approximately the same. The position of the peak does not vary with the parameters, whereas its size decreases with *α*, is weakly modulated by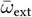, and does not change with Δ*τ*.

### 2.3 Sinusoidal forcing

We exposed the fish to an external flow rate of the form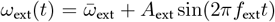, with 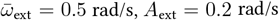and *α* = 3 cm^*−*1^ and for 5 logarithmically spaced values of the forcing frequency *f*_ext_. (Fig. S6A). Each current lasted for 30 s and was repeated twice for each frequency.

For low enough frequencies, the swimming speed *V* (*t*) follows the modulation of the target *ω*_ext_(*t*)*/α*, but one also observes the characteristic fluctuations superposed on top of it. When the input frequency is close to that of the fluctuations, the fluctuating output becomes phase-locked to the input. We can get the predicted response from the delayed linear model by considering 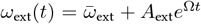 and looking for solutions oscillating at the same frequency *Ω* as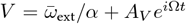. By substituting it into Eq. 10 one gets:

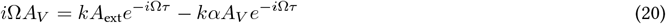

By rearranging we obtain the transfer function *H* between *ω*_ext_ and *αV* :

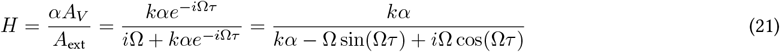

The absolute value of *H* gives us the ratio between the amplitudes of output and input:

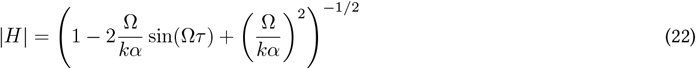

Whereas the phase of *H* gives us the phase lag between input and output:

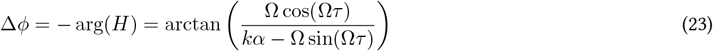

These results hold in the regime where the system is stable (*µ < π/*2), otherwise for the linear model one expects a diverging trajectory with no convergence to a steady state. The presence of noise makes it hard to quantify the amplitude of the speed oscillations, but we can easily compute the time delay Δ*t* between *ω*_ext_(*t*) and *V* (*t*) by calculating their cross-correlation and finding the time at which it has a maximum. Then the phase lag is simply given by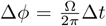. This way we measured the phase lag for different values of the forcing frequency.

We then fit this data with the expression that we derived from the model (Eq. 23), which depends on the two independent parameters *τ* and *k*. We find that the data are well fit by the model (Fig. S6B). From the fit, we extracted an estimate of the delay *τ* = 150 *±* 10 ms, which matches the value estimated from the impulse response function measurement, and an estimate of the responsiveness *k* resulting in a dimensionless gain *µ* = 1 *±* 0.2.

### 2.4 Increasing the feedback delay

To further test the validity of the delay model to describe the system, we manipulated the feedback delay by simply waiting for some extra time Δ*τ* before updating the visual environment, rather than doing immediately. *V* (*t*) being the swimming speed computed at time *t*, the net flow rate transmitted to the projector at time *t* becomes *ω*(*t*) = *ω*_ext_(*t*) *− αV* (*t −* Δ*τ*). We presented the fish with constant currents of 30 s duration with *ω*_ext_ = 0.5 rad/s and *α* = 1.5 cm^*−*1^ in randomized trials with either Δ*τ* = 0 or 5 logarithmically spaced values of Δ*τ >* 0, with two distinct trials for each value of the additional feedback delay Δ*τ*.

From the theoretical analysis of the linear delay equation, we expect that an increase of the delay *τ* should eventually move the system to the unstable regime, leading to diverging oscillations. As discussed, we expect the linear model to break down and possibly to observe limit cycle oscillations because of the presence of nonlinearities. In particular we observed that fish can only swim in a limited range of speed, therefore we might expect the acceleration to decrease as the boundaries of this physiological range are approached, regardless of the visual flow.

We observed that for a large enough value of the delay, the oscillations become more regular and are confined within the range of accessible speed values, suggestive of limit cycle oscillations (Fig. S6C). We also noted that the frequency of the fluctuations decreases with the delay. This behavior is expected from the linear delay model. Reverting back to actual physical dimensions for the solution of Eq. 16, we obtain the predicted oscillation frequency:

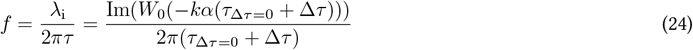

We expect this relationship to hold in the stable spiral regime, whereas in the unstable regime the oscillation frequency is determined by the nonlinearity and might deviate from Eq. 24. We estimated the oscillation frequency from the experimental data by computing the average spectrogram of 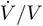 and extracting its peak frequency. We fit Eq. 24 to the experimental data and found that it describes it well (Fig. S6D). The fitting procedure is underdetermined, resulting in large relative uncertainties of the best fit values of *τ*_Δ*τ*=0_ and *k*. We can still fix the measured value of *τ*_Δ*τ*=0_ *≈* 150 ms in the fit and get an estimate of *k*, resulting in a dimensionless gain *µ*_Δ*τ*=0_ = 1.3 *±* 0.2. For a linear system with a fixed responsiveness *k* we would expect Eq. 24 to break down for *kατ > π/*2, corresponding to Δ*τ >* (*π/*(2*g*_Δ*τ*=0_) *−* 1)*τ*_Δ*τ*=0_ *≈* 0.04 ms, as one transitions to the unstable regime. Yet, we find the Eq. 24 seems to capture the frequency of the oscillations also for large values of the delay. We will see that this kind of dependence actually results from the combined effect of noise and nonlinearity.

### 2.5 Responsiveness estimation via linear regression

We estimated the value of the responsiveness *k* for different parameters by analyzing the relationship between 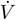 and *ω*(*t − τ*) in the measured data. To do so, we considered the value of *τ* extracted from the impulse response function. We performed a linear regression of the acceleration on the perceived optic flow rate separately for each fish and each condition, using experiments performed under constant currents. We convolved the optic flow rate with a Gaussian (standard deviation 20 ms) to account for the effect of the response kernel. Moreover, we denoised the acceleration with another Gaussian convolution (standard deviation 30 ms). To reduce the effect of outliers, we performed a robust regression using the Theil-Sen estimator [20, 21].

We randomly sampled a fraction of the data points to make the computation faster, and excluded outliers by ignoring flow rate values below the 0.1 quantile and above the 0.9 quantile. This method gave us estimates of *k* from which we computed corresponding estimates of the dimensionless gain *µ* = *kατ*. For the experiments with increased feedback delay we used *τ* = *τ*_0_ + Δ*τ*, where Δ*τ* is the artificially added delay.

### 2.6 Open-loop perturbations

We performed experiments with a constant current *ω*_ext_ in closed-loop and then suddenly switched off the feedback and imposed a current directed along the fish heading direction with a flow rate *ω*. We repeated this sequence for different values of the perturbation flow rate *ω*, both forward *ω >* 0 and backwards *ω <* 0. We presented the different perturbations in a randomized order for a duration of 0.5 s with a variable closed-loop time of 8–8.5 s in-between perturbations.

For a given baseline closed-loop condition (*ω*_ext_, *α*) we repeatedly presented the different values of *ω* over at least 10 trials separated by pauses of 1 minute with no external current Then, we repeated the experiments for different values of *ω*_ext_ and *α* to study the effect of the baseline condition. We observed that, in response to the perturbation, the fish vary their swimming speed approximately exponentially after a certain delay (Fig. S8A,B). We thus fit the trajectories with the following functional form:

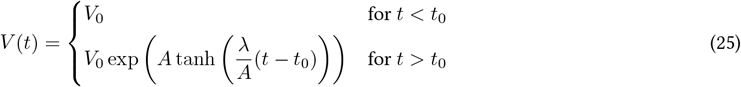

We have included a saturation in the form of an hyperbolic tangent to account for the fact that the fish speed is physiologically bounded. We excluded the trials where the fish were not swimming at the beginning of the perturbation. We used the remaining data to extract the best fit estimates of *λ, A, t*_0_ and *V*_0_. We found the acceleration rate *λ* to be approximately independent of the closed-loop conditions, including the target speed *V* ^*\**^ just before the perturbation (Fig. S8D). This would not be expected for a constant acceleration independent of the current swimming speed *V* and it further supports the idea of exponential dynamics.

Then, for each fish, we computed the median of the trajectories of *V* (*t*) across all trials and for all conditions, in order to reduce the effect of outliers and capture the typical response. We fit the response again to extract the acceleration rate *λ* for each fish as a function of the optic flow rate *ω*. We also noticed that the delay *t*_0_ is compatible with our measured value of *τ* from the impulse response function, but actually shows a dependence on the flow rate. We find that the delay before accelerating in response to forward flows is smaller than that before deceleration for backward flows, but it remains in the range of *≈* 100–200 ms (Fig. S8C). For each fish we fit the dependence of *λ* on *ω* with the following functional form:

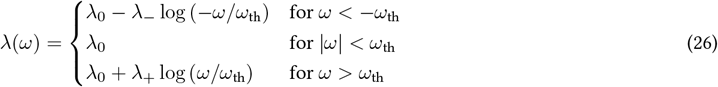

We extracted estimates of the fitting parameters *λ*_0_, *λ*^*−*^, *λ*_+_ and *ω*_th_. We found that the response of individual fish is well described by Eq. 26 and the parameters have comparable values across fish (Fig. S8E). We find that the threshold is *ω*_th_ = 0.03 *±* 0.01 rad/s and the baseline deceleration rate is *λ*_0_ = *−*0.7 *±* 0.6 s^*−*1^. We also noted an asymmetry in the response to forward and backward flows in some of the fish. We do not expect the response to optic flow values below threshold to be a good description of the behavior of fish that are stabilizing a current in closed-loop. This is because in such situation the flow rate fluctuates quickly between positive and negative values without remaining close to zero, and we think that the deceleration of fish in the open-loop perturbations is triggered by the sustained absence of flow. Because of this, for our model we considered a logarithmic response without a threshold, by shifting the logarithm to the origin:

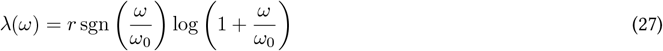

We have also made the approximation of a symmetric response for positive and negative flow rates. To extract the values of *r* and *ω*_0_ we again fit the acceleration rates for each fish but excluded the values below the threshold |*ω*| *< ω*_th_ and symmetrized the response for positive and negative *ω*. By combining the results for the different fish we obtained the estimates *r* = 1.6 *±*0.3 s^*−*1^ and *ω*_0_ = 0.07 *±* 0.03 rad/s.

### 2.7 Noise amplitude estimation

We tried to estimate the functional form of the deterministic and stochastic terms in the dynamics of the swimming speed. To do so we considered a generic stochastic differential equation for the dynamics of *V* :

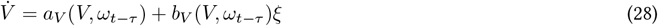

Where *ξ* is delta-correlated Gaussian white noise. In general both the drift term *a*_*V*_ and the noise amplitude *b*_*V*_ can depend on the perceived optic flow rate and on the swimming speed itself. The equation can be integrated over a small time interval Δ*t* to get:

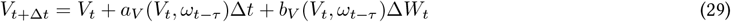

Where we considered Itô’s interpretation of the noise term and Δ*W*_*t*_ is the increment of a the Wiener process *W*_*t*_, for which ⟨Δ*W*_*t*_⟩ = 0 and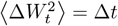. Then we can estimate the drift term and noise amplitude as the average change and variance of *V* :

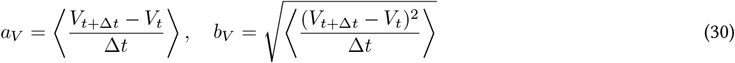

Similarly by considering the equation for log(*V*) we get:

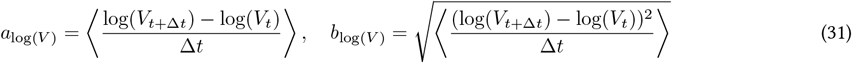

We computed these estimates for the experiments with constant external flow rate in different conditions. We used the estimate of the delay obtained from the impulse response function. We binned the values of the perceived optic flow rate, choosing an adaptive bin width such that each bin contained the same number of data points, then we evaluated the median and median absolute deviation of the computed finite differences to get estimates of drift and noise amplitude as functions of the perceived optic flow rate (Fig. S9). We found that the noise amplitude for *V* shows a dependence on *α, ω*_ext_ and *ω*| _*t − τ*_. In contrast, when we evaluate the noise amplitude for log(*V*), the curves collapse and the result is independent of the parameters, yielding an estimate *σ* = *b*_log(*V*)_ = 0.9 *±* 0.2 s^*−*1*/*2^. This suggests that the noise in the dynamics of log(*V*) is additive, and does not depend on *V* or *ω*| _*t − τ*_. We observe a similar collapse for the drift term, where estimates for different values of *α* and *ω*_ext_ align together on the same function of the perceived optic flow. This collapse confirms our observation that the dynamics of *V* is exponential, therefore *a*_*V*_ depends on the target speed itself, whereas such a dependence vanishes in *a*_log(*V*)_. As expected the drift is positive for *ω*| _*t − τ*_ *>* 0 and negative for *ω*| _*t − τ*_ *<* 0. The data collapse for the drift is quite clear except for different values of *ω*_ext_ for *ω*| _*t − τ*_ *>* 0, where it looks like the different curves actually move away from each other as one moves from the linear to the logarithmic estimates. This suggests an additional dependence of the acceleration on the external flow rate *ω*_ext_ that is not captured by our model.

### 2.8 Stochastic nonlinear delay equation analysis

Here we study the solutions to the stochastic nonlinear delay equation:

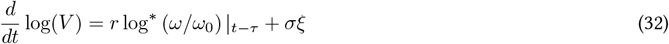

Where *ξ* is a delta-correlated Gaussian white noise. We consider the case of a slowly varying *ω*_ext_. Expanding the derivative of the logarithm one obtains a nonlinear equation for *V* with a dependence both on *V*| _*t*_ because of the multiplicative response and on *V*| _*t − τ*_ because of the feedback. This equation can be integrated numerically for a given realization of the noise, but it is hard to study analytically. If the speed remains of the order of the target *V* ^*\**^ we can simplify the analysis by considering the the approximation of small oscillations |*V/V* ^*\**^ *−* 1| *≪* 1, leading to:

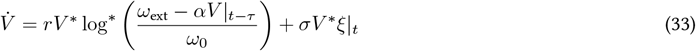

Then we can make the change of variable *x*(*t*^*′*^) = (*ω*_ext_ *− αV* (*t*))*/ω*_0_ with *t*^*′*^ = *t/τ* leading to:

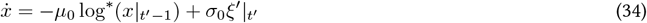

where we defined a dimensionless gain *µ*_0_ = *rτω*_ext_*/ω*_0_ *σ*_0_, a dimensionless noise amplitude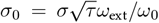, and 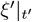 is a Gaussian white noise such that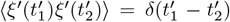. The factor of 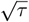 appears upon rescaling the noise term as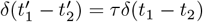.

#### Deterministic case

We first study the deterministic case, *σ*_0_ = 0, leading to the nonlinear delay equation:

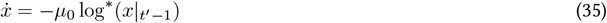

For *x ≪* 1 one can linearize the response, log^*\**^(*x*) *≃ x*, yielding a linear delay equation with dimensionless gain *µ*_0_. In this case, the behavior of the solutions close to the fixed point *x* = 0 is determined by the value of *µ*_0_, as we detailed in the analysis of the linear delay equation. For *µ*_0_ *> π/*2 the fixed point becomes unstable and one gets diverging oscillations. However, in the case of the nonlinear equation Eq. 35, the response is a sublinear function of *x* and therefore we expect the solution to settle on a limit cycle. To estimate the amplitude of the limit cycle oscillations, we extended the ideas derived from the analysis of the linear delay equation. For a certain value of 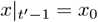 the system evolves locally with a rate of change *−µ*_0_ log^*\**^(*x*_0_) which would equivalently arise from a linear system with a gain *µ*_0_ log^*\**^(*x*_0_)*/x*_0_ (Fig. S10A). With this little trick, we can locally describe the nonlinear system as a linear one. Then we expect the solutions to diverge to infinity where *µ*_0_ log^*\**^(*x*_0_)*/x*_0_ *> π/*2 and converge to zero where *µ*_0_ log^*\**^(*x*_0_)*/x*_0_ *< π/*2. We thus expect the amplitude of the limit cycle oscillations to roughly correspond to a value of *x*_0_ at the transition between stability and instability, i.e. *µ*_0_ log^*\**^(*x*_0_)*/x*_0_ *≃ π/*2. We can define the effective amplitude of the limit cycle oscillations *A*_lc_ with the relationship *µ*_0_ log(1 + *A*_lc_)*/A*_lc_ = *π/*2. More generally we can define the effective amplitude *A*_e_ corresponding to the effective gain *µ*_e_ with the relationship *µ*_0_ log(1 + *A*_e_)*/A*_e_ = *µ*_e_. Then we can solve for *A*_e_ in terms of the Lambert function to get:

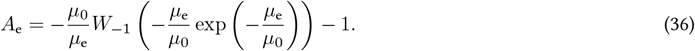

For the effective limit cycle amplitude one finds:

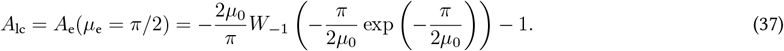

This expression equals zero for *µ*_0_ *< π/*2 and grows linearly as *A*_lc_ *≃* 4*/π*(*µ*_0_ *−π/*2) close to the bifurcation for *µ*_0_ *> π/*2. For large values of the bifurcation parameter *µ*_0_ *≫ π/*2 the amplitude grows as *A*_lc_ *≃ µ*_0_ log(*µ*_0_). These scalings can be obtained by using the fact that *W*_*−*1_(*x*) *≃* log(*−x*) for *x →* 0^*−*^ and 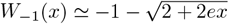for 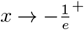.

We note that the scaling close to the bifurcation differs from that of a canonical Hopf bifurcation. The reason behind this can be seen by performing a Taylor expansion of the nonlinearity. For *x ≪* 1, one gets 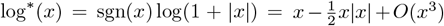 which has a quadratic term in *x*, whereas the normal form for the Hopf bifurcation has a cubic nonlinearity.

We have derived an expression for the effective amplitude of the limit cycle oscillations *A*_lc_, which we expect to set the actual scale of the oscillations. We ran simulations of Eq. 35 for various values of *µ*_0_ and extracted the oscillation amplitude at steady state by looking at the standard deviation of the *x* values (Fig. S10B). By comparing these measurements with *A*_lc_ we found that they are well described by std_lc_ = *c*_lc_*A*_lc_ with *c*_lc_ *≈* 0.9.

#### Effect of the noise

The effect of the noise term is to drive the solutions away from the attractors of the deterministic equation. In doing so, it can amplify some modes of the deterministic dynamics and lead to noise-induced oscillations. To understand the scale of the fluctuctations induced by the noise we start by considering the case of a Ornstein-Uhlenbeck process [22]:

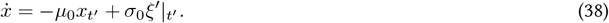

This is a well known stochastic process, corresponding to an overdamped harmonic oscillator driven by noise. It can provide a good description of the system in the limit where both *µ*_0_ and *σ*_0_ are small and the deterministic dynamics have a stable fixed point with a long relaxation time and the noise induces small amplitude fluctuations around it, so that we can approximately linearize the logarithmic nonlinearity and ignore the effect of the delay. By solving the corresponding Fokker-Planck equation, one finds that the steady-state probability distribution of *x* is a Gaussian with zero mean and standard deviation std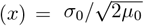. We see that the fluctuation amplitude thus grows with the noise amplitude *σ*_0_ and decreases with the strength of the restoring force *µ*_0_. We have found that the proper limit in which the fluctuation amplitude is small std(*x*) *≪* 1 is 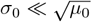.

If we relax the condition *σ*_0_ *≪* 1, we cannot linearize the logarithm anymore, but we can still use the local linear approximation. We obtain an effective gain *µ*_n_ for the effective amplitude *A*_n_ induced by the noise as *µ*_0_ log(1 + *A*_n_)*/A*_n_ = *µ*_n_. Then we expect the standard deviation of the fluctuations to be determined by this effective gain as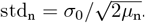. The effective amplitude should be of the same order as the standard deviation, so we can write std_n_ = *c*_n_*A*_n_ with *c*_n_ *≃* 1. By equating the effective gain from the two relationships, we obtain an implicit relation for the effective amplitude of the noise-induced fluctuations:

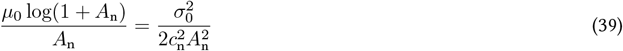

This relation has a single solution in *A*_n_ which can be easily evaluated numerically for a given choice of the parameter values. We have run simulations of Eq. 34 for various values of *µ*_0_ and *σ*_0_ and found that the standard deviation of the *x* values is well described by Eq. 39 for small values of *µ*_0_ (Fig. S10B). We varied *c*_n_ to find the value for which the theoretical estimate best matches the simulation results and found *c*_n_ *≈* 0.7.

For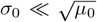, one can linearize the logarithm in Eq. 39 and recover the expected result for a linear system with gain *µ*_0_. In the opposite limit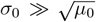, we can approximately write 1 + *A*_n_ *≃ A*_n_ inside the logarithm and solve in terms of the Lambert function to get *A*_n_ = *s/W*_0_(*s*) with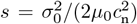. In this limit, the expression can be further simplified by considering the asymptotic approximation of the Lambert function: *W*_0_(*s*) *≃* log(*s*) *−* log(log(*s*)).

For large values of *µ*_0_ we expect the amplitude of the solutions to be dominated by the limit cycle oscillations and follow std_lc_ = *c*_lc_*A*_lc_, which is indeed what we observe in the simulations (Fig. S10B).

We found theoretical estimates of the fluctuation amplitude in the limits where the solution is either dominated by the noise or by the limit cycle, in general we expect the fluctuation amplitude of the solutions to be the sum of the two contributions:

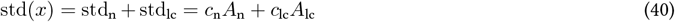

Where *A*_n_ is given by Eq. 39 and *A*_lc_ is given by Eq. 37. For *µ*_0_ *< π/*2, *A*_lc_ = 0 and only *A*_n_ contributes to the fluctuation amplitude. From Eq. 39 we see that *A*_n_ is a continuous monotonically increasing function of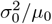. Then for *µ*_0_ *> π/*2 one has also the contribution of *A*_lc_, which is a continuous monotonically increasing function of *µ*_0_. We note that *A*_lc_ has a kink (a discontinuity of the first derivative) at the bifurcation point *µ*_0_ = *π/*2, which is reflected as a kink in the total fluctuation amplitude as it is a linear combination of *A*_n_ and *A*_lc_.

When comparing this expression with the simulation results we find that it describes them quite well even in the intermediate regime (Fig. S10D). We find a discrepancy for small *σ*_0_ close to the bifurcation *µ*_0_ = *π/*2 where the theory underestimates the amplitude and for large *σ*_0_ and small *µ*_0_ where we observe some variability in the amplitude estimates because of the limited simulation time. Therefore we find that there is a smooth transition between noise-induced fluctuations and limit cycle oscillations and we can define the boundary between the two regimes in the phase space defined by (*µ*_0_, *σ*_0_) as the curve for which std_n_ = std_lc_.

Now that we have computed the standard deviation of the fluctuations, we can try to extract an effective gain *µ*_e_ by going back to the effective amplitude *A*_e_. We have seen that the scaling between effective amplitude and standard deviation is slightly different in the two regimes. We can linearly interpolate between the two expressions by weighing them with the relative amplitude of the two contributions, yielding *A*_e_ = std */c*_i_ with *c*_i_ = (std_n_ *c*_n_ + std_lc_ *c*_lc_)*/*(std_n_ + std_lc_). Subtituting the definitions back in we get:

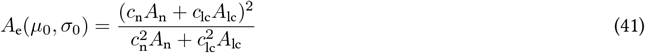

Note that for *c*_n_ = *c*_lc_ this expression simplifies to *A*_e_ = *A*_n_ + *A*_lc_. Then, the effective gain is simply given by *µ*_e_ = *µ*_0_ log(1 + *A*_e_)*/A*_e_ (Fig. S10C). From this effective gain we can also estimate the characteristic frequency of the oscillations using the prediction of the linear model: *f* ^*′*^ = Im(*W*_0_(*−µ*_e_))*/*(2*π*). Thus we find that there is a sharp transition to oscillatory behavior corresponding to *µ*_e_ = 1*/e* and then, as *µ*_0_ increases, the frequency of the oscillations grows smoothly from zero to 1*/*4 as the limit cycle becomes dominant over the noise. We have extracted the characteristic oscillation frequency from the simulation results and found that it matches our theoretical prediction (Fig. S10E), except for a narrow region around the transition to oscillatory behavior at *µ*_e_ = 1*/e*, where the characteristic timescale diverges and it cannot be extract from a finite simulation time.

#### Dependence on the physical variables

We can go back to the original variables by appropriately rescaling our results. We find that *V* fluctuates around *v*_*c*_*/α* with a standard deviation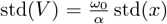. The relative fluctuation amplitude 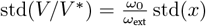 is thus independent of the feedback gain *α*.

For the characteristic frequency we have to divide by *τ* to go back to our original time dimensions *f* = *f* ^*′*^*/τ*.

As one varies the external flow rate *ω*_ext_, both *µ*_0_ and *σ*_0_ vary as they are proportional to this parameter. If one looks at the phase space (*µ*_0_, *σ*_0_) on a double logarithmic scale this corresponds to moving along a line with slope 1. Instead if one varies the delay *τ, µ*_0_ increases linearly whereas *σ*_0_ shows a square root dependence. This correspond to moving along a line with slope 1*/*2 in the same space. Varying *ω*_0_ would have the same effect as varying *ω*_ext_, while varying *r* and *σ* respectively results in horizontal and vertical displacements in the parameter space.

Thus by increasing *ω*_ext_ or *τ* we can move from a regime with no oscillation to one where the system fluctuates with a characteristic frequency, and one can transition from a regime where oscillations that are dominated by the noise to one where they are dominated by the deterministic dynamics.

Since *A*_n_ only depends on the parameter combination 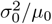, it grows with *ω*_ext_ but does not depend on *τ*. On the other hand, *A*_lc_ depends on *µ*_0_ and grows both with *ω*_ext_ and *τ*. As a result, the fluctuation amplitude always increases with *ω*_ext_ and increases with *τ* for *µ*_0_ *> π/*2 whereas it is independent of *τ* for *µ*_0_ *< π/*2.

We can substitute the definitions of *ω*_ext_ and *τ* to get the scalings:

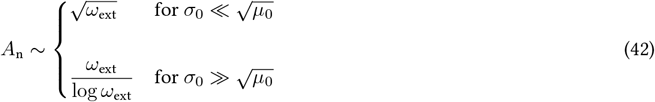

And:

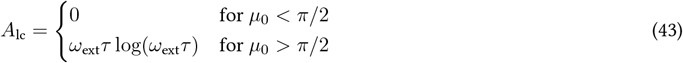

Thus the relative fluctuation amplitude std(*V/V* ^*\**^) *∼* (*A*_n_ +*A*_lc_)*/ω*_ext_ diverges as 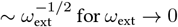 and then shows a weak (sublinear) dependence on *ω*_ext_, decreasing as 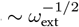 and possibly *∼* 1*/* log(*ω*_ext_) for *µ*_0_ *< π/*2 and eventually increasing as *∼* log(*ω*_ext_) for *µ*_0_ *> π/*2. In contrast, as one varies *τ*, the relative fluctuation amplitude remains constant for *µ*_0_ *< π/*2 and then increases with a stronger functional dependence as *∼ τ* log(*τ*) for *µ*_0_ *> π/*2.

#### Comparison with experimental data

We used the estimated values of (*τ, σ, r, ω*_0_) to derive theoretical estimates of the relative fluctuation amplitude std(*V/V* ^*\**^), characteristic frequency *f*, and effective gain *µ*_e_, for different values of *α, ω*_ext_, and Δ*τ*. As these quantities do not have a continuous derivative as a function of the parameters, we could not simply propagate the uncertainties using the Taylor expansion. We decided to use a Monte Carlo approach, by repeatedly sampling a multivariate Gaussian distribution with the mean and covariance that match the estimated distribution of (*τ, σ, r, ω*_0_). We took into account the strong correlation between *r* and *ω*_0_, estimated to be *≈* 0.95 from the fitting procedure of Eq. 27. Then we computed the theoretical estimates for each of the sampled set of parameters values, and we extracted the corresponding confidence interval by evaluating the mean and standard deviation of such estimates.

### 2.9 Numerical simulations

To test the nonlinear model and the limitations of the effective linear theory, we performed numerical simulations of Eq. 32. As the theory was developed in the limit of small oscillations we expect it to break down for large fluctuations. We also included in the simulations some features of the experimental system that we ignored in the model. A first modification consists in taking into account the finite width of the response kernel: instead of considering *ω*| _*t − τ*_ in the response, we plug in the convolution *G * ω* where we approximate the response kernel *G* as a Gaussian with mean 150 ms and standard deviation 20 ms. For the remaining parameters (*r, ω*_0_, *σ*) we considered the estimates that we obtained from the experiments. Moreover we included the effect of saturation at small and large values of the swimming speed. We did it by setting the deterministic part of the response to zero if it was positive when *V > V*_max_ = 0.9 cm/s or negative when *V < V*_min_ = 0.09 cm/s. The resulting equation is:

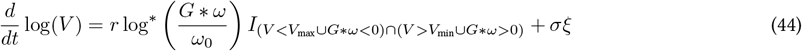

Where *I* denotes the indicator function of the set of values where the deterministic part is not suppressed. We simulated the equation using the Euler method with a time increment of 10 ms. We considered constant currents of 15 minutes duration for different parameters values and then analyzed the resulting simulated data as we did for the experimental data. We found that the resulting trajectories quantitatively match the experimental data and show similar deviation from the theory due to speed saturation at large values of *V* ^*\**^ (Fig. S11).

## 3 Neural activity

### 3.1 Light-sheet microscope with confocal slit detection

We recorded brain activity using a custom-made light-sheet microscope adapted from the one described in [23]. A laser emitting blue light at *≈* 488 nm is used to excite the fluorescence of calcium sensor in the neurons. Along its optic path, the beam is focused using scanning and tube lenses onto two galvanometric mirrors that can rotate around perpendicular axes, resulting in horizontal and vertical translations at the sample. The laser beam is focused by an illumination objective (Zeiss EC Plan-Neofluar 5x/0.16) mounted on the side of the water tank, through a glass window and into the fish brain. At the sample, the focused beam displays a gaussian profile with a thickness of *≈* 4.5 µm (full width at half maximum) and a power of *≈* 150 µW. The green light emitted by the calcium sensor is collected by the detection objective placed above the fish and focused on the sensor of a camera (Hamamatsu ORCA-Flash4.0 V3 C13440-20CU, 2048 *×* 2048 pixels of size 6.5 *×* 6.5 µm). We used a Thorlabs TL20X-MPL water dipping objective associated with a tube lens of focal length 100 mm, resulting in a magnification of 10x. In-between the camera and the detection objective, we inserted a notch filter (Thorlabs NF488-15) to block the light from the laser, a band-pass filter (Semrock FF01-520/35-25) to block the light from the projector, and a short-pass filter (Semrock FF01-720/SP-25) to block the IR light from the LED. We also added a magenta dichroic filter (Thorlabs FD1M, mounted at an angle of *≈* 30^*°*^) just after the projector to remove green light from the visual stimulus so that it does not interfere with the fluorescence signal. To prevent the laser light from impinging directly onto the eye of the fish, we added a small metal shield on the optical path of the beam, and mounted an additional camera and mirror to position it properly before each experiment. To acquire an image of the fluorescence in one thin layer of the brain, we scan the laser beam horizontally across the sample by rotating one of the mirrors. Layers at different heights in the brain are successively imaged by changing the height of the beam using the other mirror while simultaneously varying the height of the detection objective using a piezo nanopositioner (piezosystem jena MIPOS 500) connected in closed-loop in order to maintain the beam in the focus plane.

One issue with this approach is that the whole camera sensor is collecting photons as the laser moves across the sample, thus scattered photons can reach pixels that do not correspond to the position from where the light was emitted. Thus part of the light emitted from the neurons ends up in the pixels corresponding to different neurons, this mixing of different signals can lead to spurious correlations in the analysis of brain activity. One way to mitigate this cross-talk is to only acquire light from a stripe of pixels that are conjugated to the position of the beam. This technique is called confocal slit detection [24] as it relies on a similar approach of optical sectioning as in standard confocal microscopy. In order to implement this method, the stripe of active pixels has to move along the sensor in sync with the movement of the beam across the sample. Our camera supports this acquisition mode and sends output signals that can be used to control the position of the beam. We programmed an FPGA (CompactRIO cRIO-9054, National Instruments) using LabVIEW to rotate the mirrors in response to the camera outputs, using NI-9263 and NI-9402 modules for analog and digital control, respectively. A graphical interface was developed in Python to access and modify the acquisition parameters. We set the stripe of simultaneously active pixels to be 20 pixels wide, equivalent to 13 µm at the sample. We decided to also move the beam vertically for each line update, this results in layers that are skewed but prevents oscillations of the piezo as it moves continuously rather than in discrete steps. At the end of a determined number of layers the vertical position is reset to the initial one. We limited the speed of this movement to prevent large acceleration that induce resonant oscillations of the piezo, and we waited for *≈* 50 ms to allow the piezo to reach the starting position before continuing with the next acquisition. During functional imaging, we acquired images with an exposure time of *≈* 15 ms per layer and a vertical distance of *≈* 8 µm between layers. Acquiring 25 different layers resulted in an acquisition rate of *≈* 2.3 brain volumes per second.

### 3.2 Imaging protocol

To perform brain recordings while fish swim in the virtual reality, we triggered the FPGA through the same Python notebook that controls the virtual reality. For the closed-loop experiments (*α >* 0, *β >* 0) we presented constant currents for three different values of the external flow rate in three randomized repetitions, with each current lasting 30 s, interspersed with pauses of 30 s. For the open-loop experiments (*α* = 0, *β* = 0), we displayed constant currents both in the forward and backward directions with 5 logarithmically spaced values of the external flow rates in three randomized repetitions, with each run lasting 20 s and pauses of 20 s in-between. For each fish we also acquired a high resolution stack of brain images with small vertical increments between layers (*≈* 1 µm) and large exposure time (*≈* 150 ms), to facilitate the mapping between different fish brains and allow for a spatial comparison of the identified neuronal populations.

### 3.3 Brain segmentation

We developed a segmentation pipeline to extract the position of the neurons from the images and their activity over time. To correct for possible motion in the horizontal place, we aligned the images in each layer by computing their cross-correlation with a chosen reference time. As we are interested in neurons that show a modulation of activity during the time of the recording, we computed the average covariance between the activity of neighboring pixels. The resulting image has a high intensity at pixels that have high local intensity correlations, therefore it is enhanced at the position of active neurons. As the calcium sensor is confined to the nuclei, each neuron appears as a spherical fluorescent object. Thus we used a blob detection algorithm that uses the Laplacian of Gaussian filter to extract the position and size of circular objects in the image [25], giving the position and radius of the nuclei. Then we extracted the fluorescence activity for each neuron by computing the mean pixel intensity for each image across all pixels belonging to that neuron.

Because of the binding kinetics of the calcium sensor, the fluorescence signal is approximately the result of a convolution between the actual calcium concentration in the neuron and a response kernel that we estimated to be an exponential decay with a characteristic time of *≈* 3 s. In order to compare the neural activity with the visual stimulus and the behavior, we deconvolved the fluorescence traces by dividing their Fourier spectrum with that of the response kernel. We further reduced the noise in the fluorescent traces by smoothing them with a Gaussian kernel with standard deviation 0.75 s. Then, to be able to compare the activity of different neurons that can express varying concentration of calcium sensor, we computed the relative fluorescence change Δ*F/F*_0_ = (*F − F*_0_)*/F*_0_, where the baseline fluorescence *F*_0_ for each neuron was taken to be the median value of *F* during the times where no visual flow was presented and the tail was not moving.

### 3.4 Brain registration

As the brains of different fish can have different sizes and can have slightly variable orientations in the agarose gel, we developed a registration pipeline to transform the brain of different fish so that they are aligned in a common space. To implement the registration we used the SimpleITK library in Python [26], which embeds the images in a physical space where transformations can be easily applied. We first mapped the high-resolution 3D stacks from all fish onto a single one. We did that by finding the optimal affine transformation between each stack and the reference one. We used correlation as similarity metric and gradient descent as optimization algorithm. Moreover, the metric was computed on a randomly sampled subset of points at each step, to increase computational efficiency. Once all stacks were mapped and resampled in the same space, we averaged the intensities together to create a reference stack from the *N* = 8 fish used in the experiments. Then, we applied a rigid transformation to the reference brain to align its left-right symmetry plane with the central plane of the image stack. We did it by mapping it onto its left-right mirrored image and then applying half of the resulting tranformation.

To map the neuron positions from each experiment to the reference stack we first mapped with a rigid transformation the low-resolution stack obtained by averaging the images recorded during the experiment onto the high-resolution stack recorded for the same fish. Then we mapped the high-resolution stack to the reference one with an affine tranformation.

### 3.5 Correlation analysis

As the animal perform speed regulation, the optic flow rate oscillates around zero with a characteristic period of *≈* 0.7 s. Since the decay time of the calcium indicator is *≈* 3 s, these fluctuation in speed and optic flow rate cannot be resolved at the neuronal level. Moreover, while the fish is swimming to stabilize itself, the visual flow depends on its swimming strength through the feedback, making it hard to disentangle the visual and motor activity in the brain. Therefore we have analyzed open-loop experiments where we present fish with forward and backward visual currents of various flow rates.

To identify all neurons whose activity is somehow related to the visual flow in an unbiased way, we computed the distance correlation [27] between the optic flow rate *ω* and the activity Δ*F/F*_0_ for each neuron across the whole recording. As opposed to the Pearson correlation coefficient, which is a measure of linear correlation, the distance correlation coefficient picks up on any kind of statistical dependence between the two signals, including nonlinear correlations. Then, to assess whether the distance correlation is statistically significant, we built a null distribution by computing the distance correlation with time-shifted versions of the signals, so that any significant dependence is lost, but their local structure is preserved. To ensure that any dependence is lost we only kept time-shifted signals whose linear correlation with the original signal was smaller than 0.25 in absolute value. We used the Benjamini-Hochberg procedure [28] to determine the set of neurons that are significantly correlated with the signal for a given false discovery rate. This ensures that the problem of multiple comparison for hypothesis testing is accounted for. We controlled the false discovery rate by setting the significance threshold to *α*_FDR_ = 10^*−*5^.

We then performed hierarchical clustering to group neurons with similar activity in an unsupervised manner, facilitating the subsequent interpretation of the results. We found large populations (*∼* 10^3^) of neurons whose activity is correlated with the flow rate independently of direction, only for forward flow and only for backward flow. We also found smaller populations (*∼* 10^2^) whose activity is anticorrelated with the flow rate.We performed the same analysis for the swimming speed *V*, and identified neuronal populations whose activity is correlated or anticorrelated with it.

This approach is useful to explore the data, but it does not provide a criterion for identifying the same populations in different fish, as the choice of clusters in hierarchical clustering is somewhat arbitrary and we have no guarantee that the clustering will be similar for different fish. Moreover, some neurons may have a mixed visual- and motor-related activity, and we would like to assign them to only one of the two groups for simplicity. To solve these issues we decided to look at the Pearson correlation of neural activity with four different signals: flow rate |*ω*|, forward flow rate *ω*_+_ = *ωH*(*ω*), backward flow rate *ω*_*−*_ = *−ωH*(*−ω*), and swimming speed *V* (*H* denotes the Heaviside step function). For each of these signals we identified neurons for which the correlation is significantly (analogous procedure as for the distance correlation) larger or smaller than zero, thus identifying eight different populations of neurons. In the cases where the same neuron was present in multiple populations, they were assigned to the category for which the correlation (or anticorrelation) was highest, thus defining non-overlapping populations.

We mapped the positions of the segmented neurons to the reference stack in order to compare their anatomical location. Because of the large number of neurons, we decided not to visualize them as individual dots but to compute a spatial density for each population using kernel density estimation. We estimated the density of a population of neurons at positions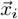 in three-dimensional space as:

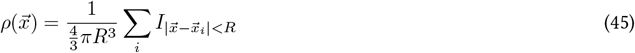

This corresponds to assigning to each unit a uniform density in a sphere of radius *R* = 5 µm surrounding its position.

To visualize this function in two dimensions, we computed the marginal distributions in two orthogonal planes by integrating in the directions perpendicular to each plane. To compare different populations 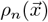 in the same image we used colors with different hues *c*_*n*_ = (*R*_*n*_, *G*_*n*_, *B*_*n*_) and computed the weighted sum of these colors with the corresponding densities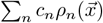. All the densities were normalized to result in a maximum pixel intensity of 1, and the weighted sum was renormalized when one of the resulting color channels exceeded 1.

To identify the brain regions associated with the neuronal populations, we reviewed the zebrafish literature and visually compared our reference stack with the Max Planck Zebrafish Brain (mapzebrain) atlas [29].

We computed tuning curves for each population as the average activity for different values of the corresponding signal. For the open-loop experiments, fish only experience a finite number of optic flow rates *ω*, so we simply took the average activity across all of the trials with the same flow rate. For the swimming speed *V*, we had to choose a finite number of bins to define the ranges for the averages, we did so by considering a bin for *V* = 0 and then positive bins with an adaptive width such that each one contained the same number of data points. We noted that the tuning curves for the populations that were negatively correlated with optic flow rate and correlated (positively and negatively) with swimming speed, showed a logarithmic dependence on the corresponding signal. We fit them as a function of the corresponding signal *x* ∈ *{*|*ω*|, *ω*_+_, *ω*_*−*_, *V}* with a logarithmic response Δ*F/F*_0_(*x*) = *A* log(*x/x*_0_)*H*(*x − x*_0_). For the populations that were positively correlated with optic flow rate we noted a mixed linear and logarithmic response. We decided to further separate them in subpopulations exhibiting either linear or logarithmic coding by comparing the correlation of each neuron’s activity with *x* and with its logarithm. We obtained distinct neuronal populations showing linear and logarithmic coding, with different spatial distributions that are consistent across fish. We computed their tuning curves and fit them either with a logarithmic response as before or a linear one of the form Δ*F/F*_0_(*x*) = *Ax*. The spatial distributions and tuning curves of all the identified populations are shown for a single fish in Fig. S12A-D. Because in open-loop experiments, fish do not swim for long periods of time, it is not always easy to extract the dependence of the neural activity on the swimming speed. We performed the same analysis on a fish swimming in closed-loop and stabilizing external currents at different speeds, finding that the tuning curves for the populations correlated with *V* are still well fit by a logarithmic response (Fig. S12E-F).

### 3.6 Effect of calcium indicator saturation

The fluorescence signal measured for a given neuron is not simply proportional to its firing rate. We thus assessed whether the observed logarithmic tuning of identified neuronal populations could arise from nonlinearities in the fluorescence response rather than reflecting the actual activity of the neurons.

The measured fluorescence *F* for a given neuron at a given time is proportional to the number of GCaMP molecules that are bound to calcium ions Ca^2+^, which is itself proportional to the total number of GCaMP molecules present in the cell and to the intracellular calcium concentration. In steady state conditions the relationship between fluorescence and calcium concentration is not linear both at high concentrations, because of saturation, and at low concentrations because of cooperative binding [30]. The fluorescence follows the Hill equation:

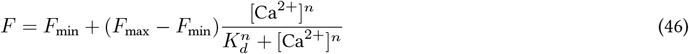

Where, for our calcium indicator (GCaMP6s), the dynamic range is *F*_max_*/F*_min_ *≃* 63, the dissociation constant is *K*_d_ *≃* 144 nM, and the Hill coefficient is *n ≃* 2.9 [2, 31]. This relationship is nonlinear but it can be approximated by a linear one in an intermediate range around the half-bound concentration *K*_d_.

Neurons at rest maintain an intracellular calcium concentrations of *≃*50 nM by actively pumping ions out as the extracellular concentration is *≃*1 mM [32,33]. Action potentials are associated with a transient increase in calcium concentration with a peak concentration of 100 nM and a decay time under *≃*100 ms [34]. During trains of action potentials the concentration reaches a steady state that grows linearly with the action potential frequency, with a slope of *≃*16 nM/Hz [34]. Physiological values of the firing rate range from less than one Hz to several tens of Hz [35, 36], corresponding to a concentration range of approximately *≃* 50–500 nM. The relationship between calcium concentration and fluorescence (Eq. 46) can be approximated as a linear one in the lower half of this range, but saturation becomes relevant as one moves in the upper half. We tried to fit the tuning curves that we measured for the identified neuronal populations by taking into account this nonlinearity to check whether it can explain the observed logarithmic responses. If we denote by *c*_0_ the calcium concentration corresponding to the baseline fluorescence *F*_0_, then by rearranging Eq. 46 (and using *F*_max_*/F*_min_ *≫* 1) we find that the relative fluorescence change corresponding to a concentration change Δ*c* is:

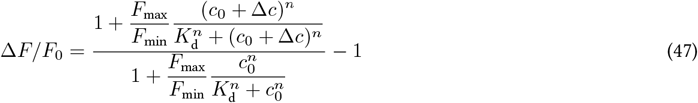

We fit the tuning curves with this expression choosing either a linear response Δ*c*(*x*) = *Ax* to the signal *x* or a logarithmic one Δ*c*(*x*) = *A* log(*x/x*_0_)*H*(*x − x*_0_). We fixed the experimental parameters of the Hill equation for GCaMP6s and left *c*_0_, *A* and *x*_0_ as free parameters. We also imposed the physiological constraint of a minimum calcium concentration of 50 nM. We found that the nonlinear response of the positively correlated populations could be explained by the saturation of the calcium indicator (even though the logarithmic fit still describes better the data), but the response of the anticorrelated populations cannot be captured by this mechanism (Fig. S13). Moreover, we identified spatially segregated neuronal populations that exhibit linear coding of optic flow rate within the same range of Δ*F/F*_0_. While we cannot exclude an effect of the calcium indicator saturation, our analysis suggests that the measured logarithmic tuning curves reflects the actual activity of the neuronal populations.

**Figure S1:**
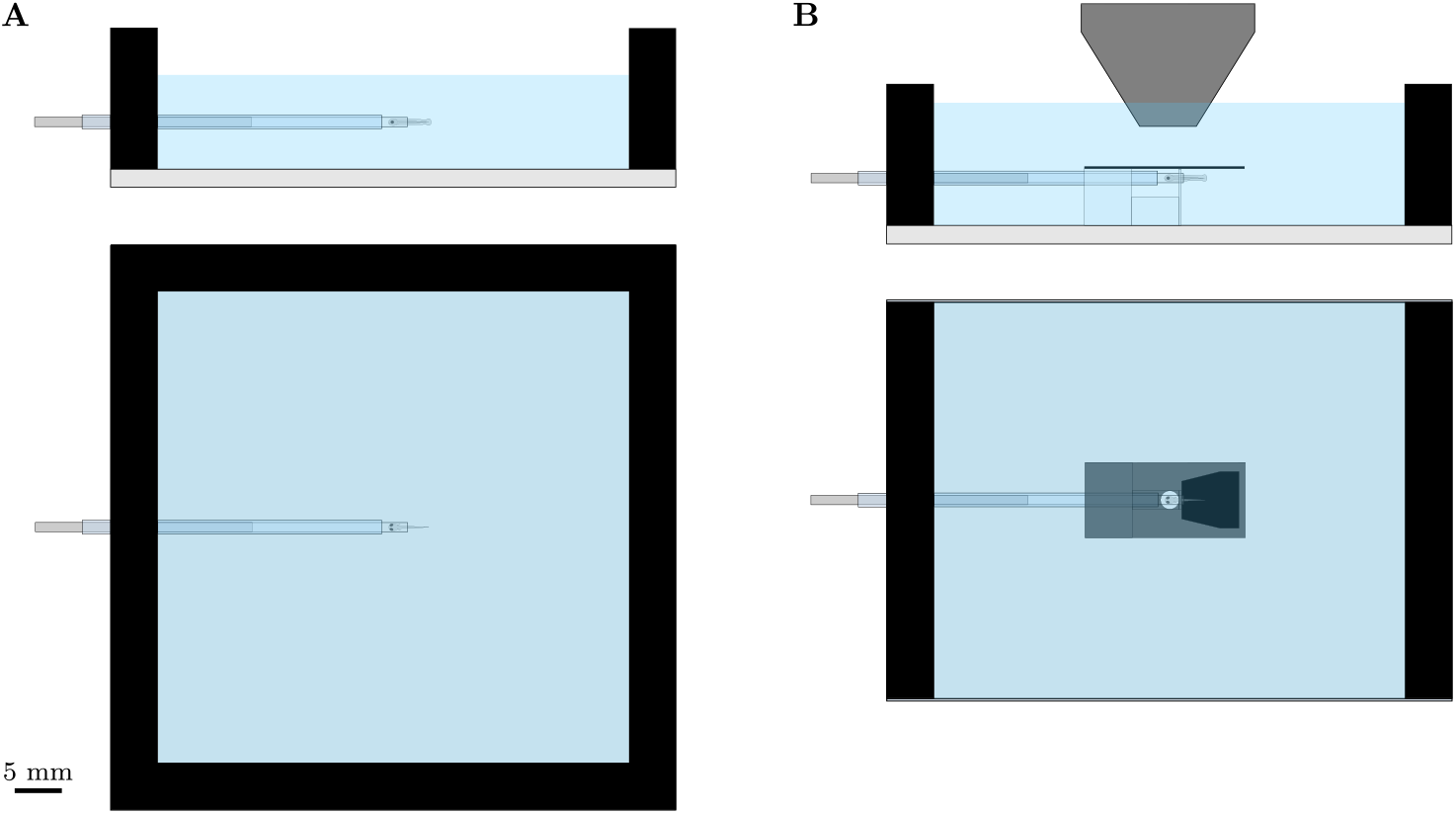
Schematic design of the water tanks. (A) Schematic of the tank used in the behavioral setup. The fish is positioned at a fixed distance from the opaque bottom using a glass capillary. (B) Schematic of the tank used in the light sheet setup, showing the detection objective above the fish. A thin mirror,placed between the fish and the objective, has a round transparent window to allow brain imaging. The projection screen on the top surface of the tank bottom, is cut out below the fish tail to enable monitoring of the tail movements with a camera. This transparent window is shown here in black. The mirror is painted black above the fish tail to facilitate the detection of the tail.

**Figure S2:**
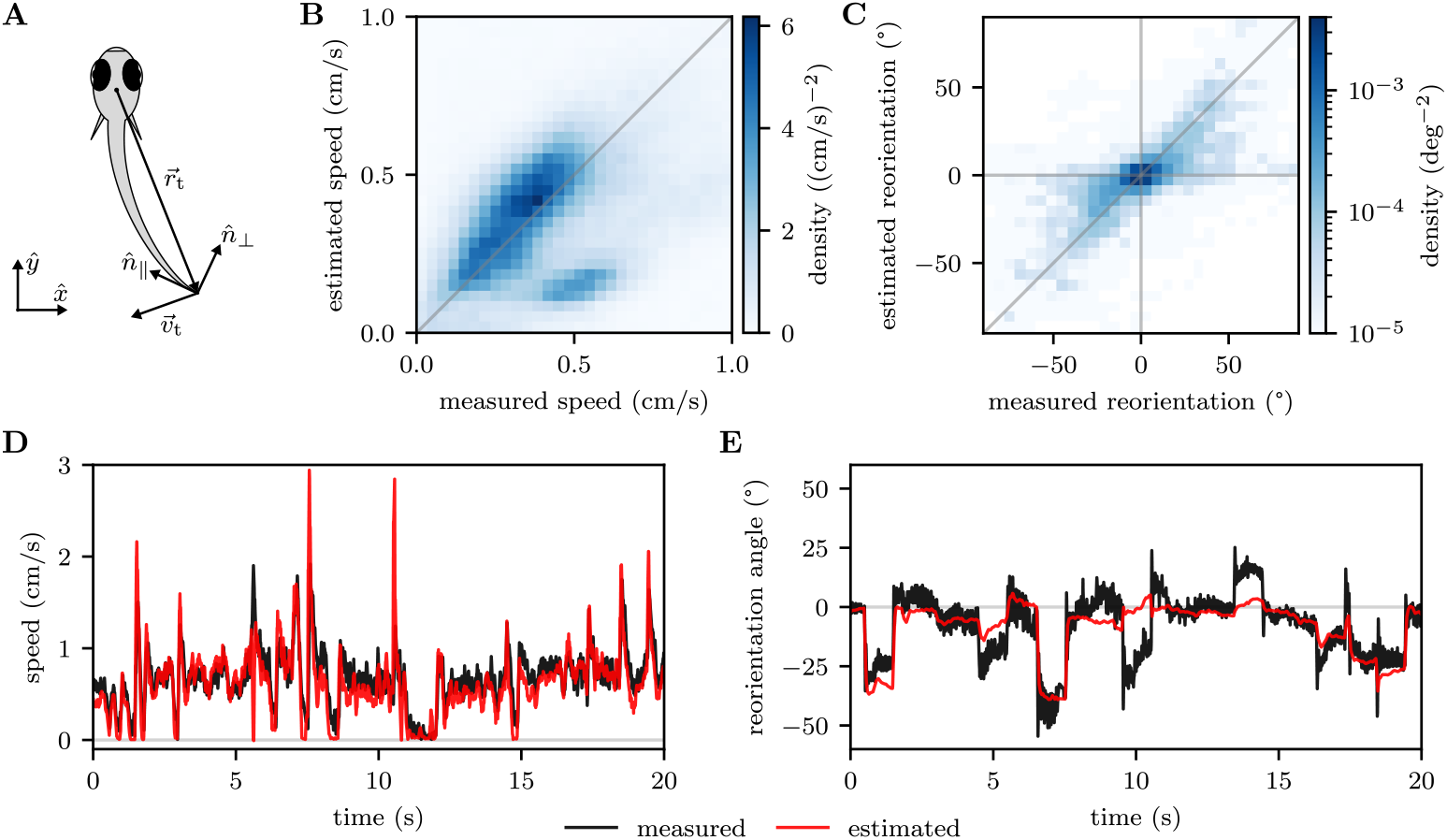
Comparison of the fictive velocity inferred from the tail movements with freely swimming data. (A) Schematic showing the relevant quantities involved in the calculation of the fictive velocity. (B) 2D Histogram (data pooled from *N* = 3 fish) showing the relationship between the measured speed *V*_fs_ and the one estimated from the tail movements *V*. Most of the data lies close to the identity line. (C) Same as (B), but for the reorientation angle *δθ*_fs_(*δt* = 1s). (D) Example traces showing the measured and estimated speed for one fish. (E) Same as (D), but for the reorientation angle.

**Figure S3:**
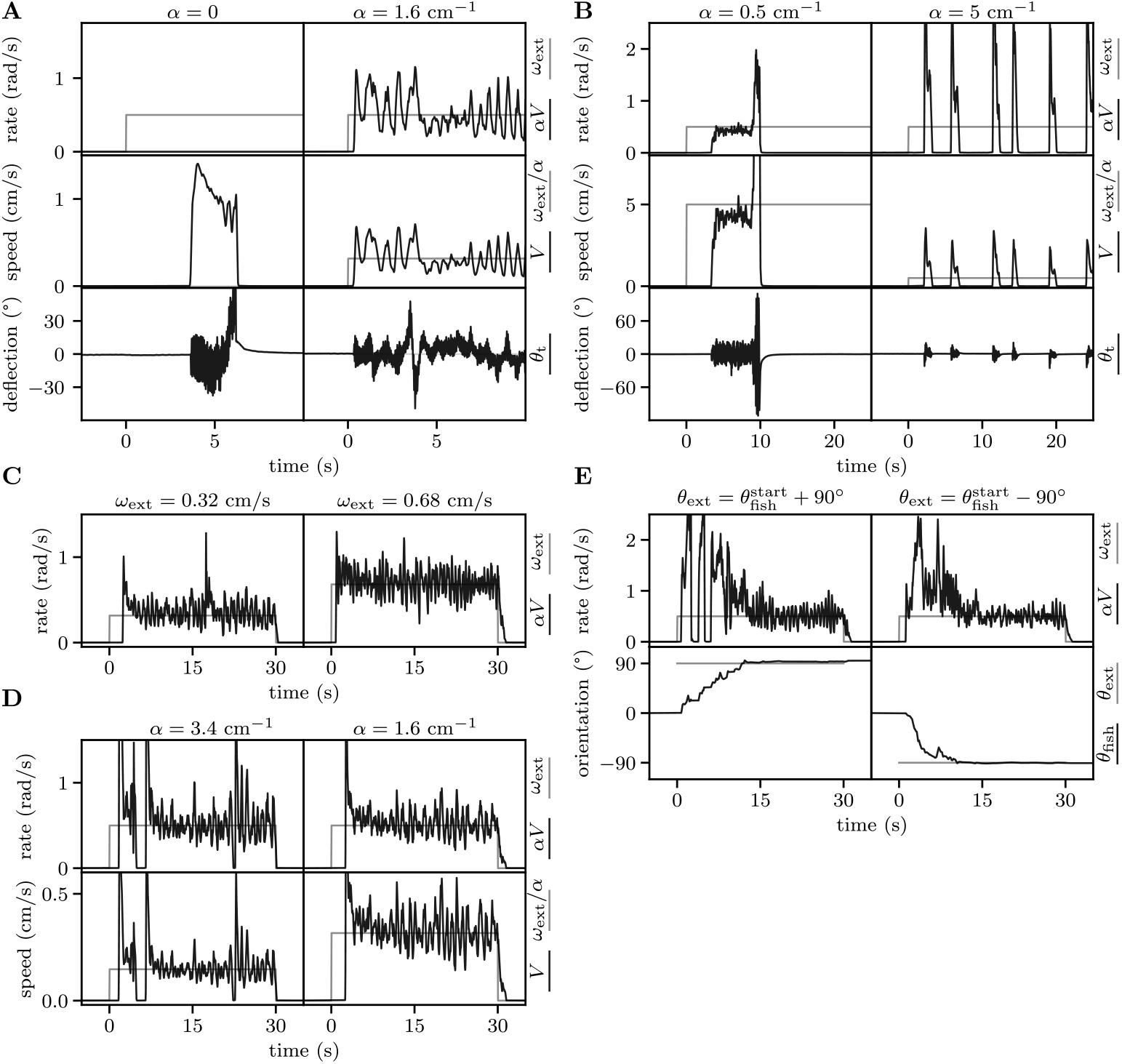
Example responses in different conditions. (A) Response of a fish to currents in open-loop (left) and closed-loop conditions (right). The tail deflection *θ*_t_ is defined in Fig. S4C. (B) Response of a fish to currents for which the target speed *V* ^*\**^ is just outside the physiological range of values that it can maintain continuously, both for large (left) and small (right) *V* ^*\**^. (C) Two example trials with different *ω*_ext_ for the same gain *α*. (D) Two example trials with different *α* for the same *ω*_ext_. (E) Two example trials with currents dragging the fish sideways from the left or from the right.

**Figure S4:**
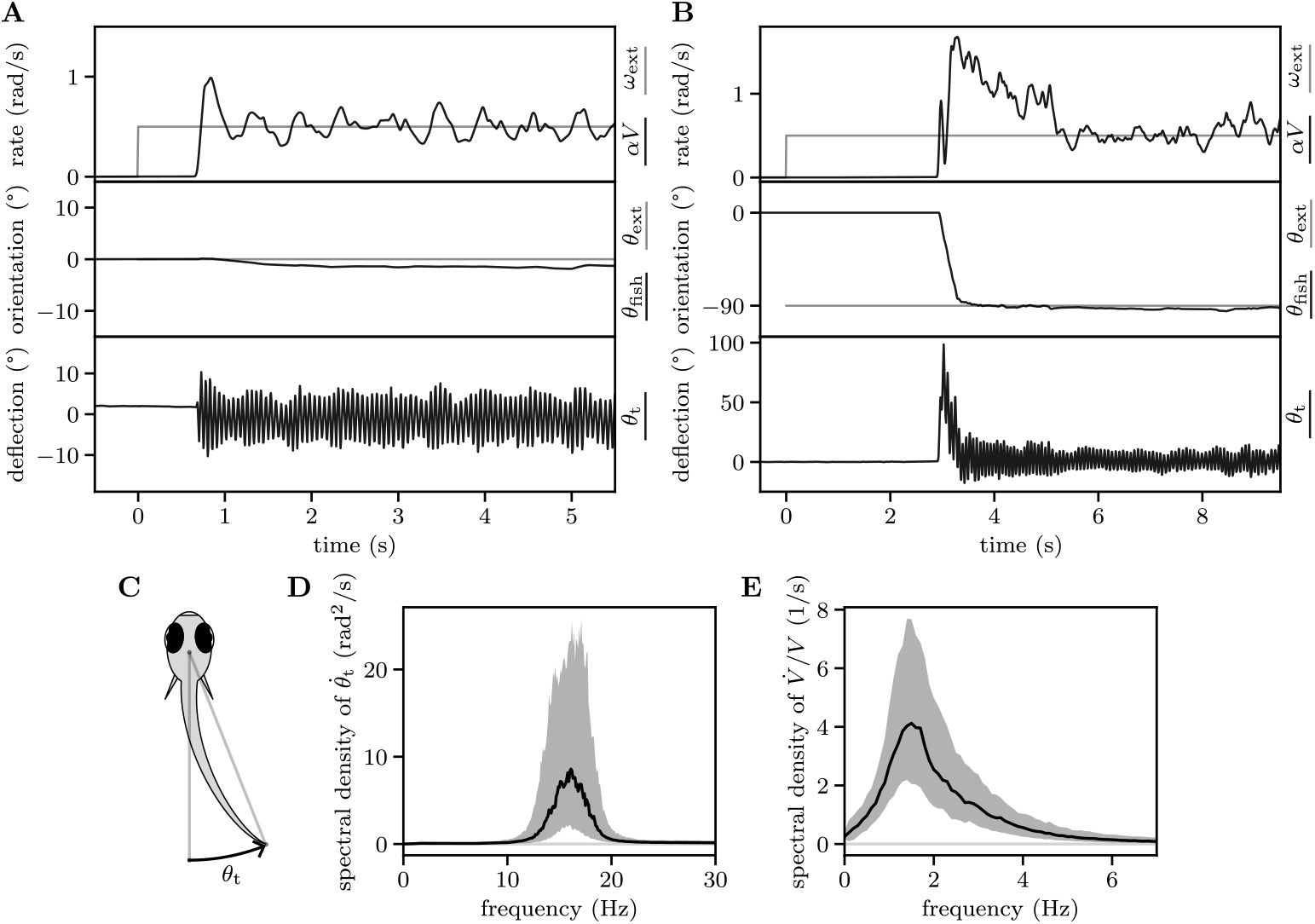
Comparison of tail deflection and speed oscillations. (A) Example response to a current in the forward direction. The fish swims by undulating its tail (bottom), it adjusts its speed to match that of the current (top) and it keeps its orientation constant to swim against the current (middle). (B) Example response analogous to (A), but with a current starting at a right angle to the fish heading direction. (C) Schematic showing the definition of the tail deflection as the angular displacement of the tail tip with respect to the head. (D) Power spectral density of the tail deflection obtained as the average spectrogram over all the experiments with constant currents (solid line and shaded area, median and interquartile range across all fish and conditions). We took the time derivative of the tail deflection *θ*_*t*_ to suppress low-frequency modulations. (E) Power spectral density of the swimming speed, computed as in (D), but in this case we considered the time derivative of log(*V*).

**Figure S5:**
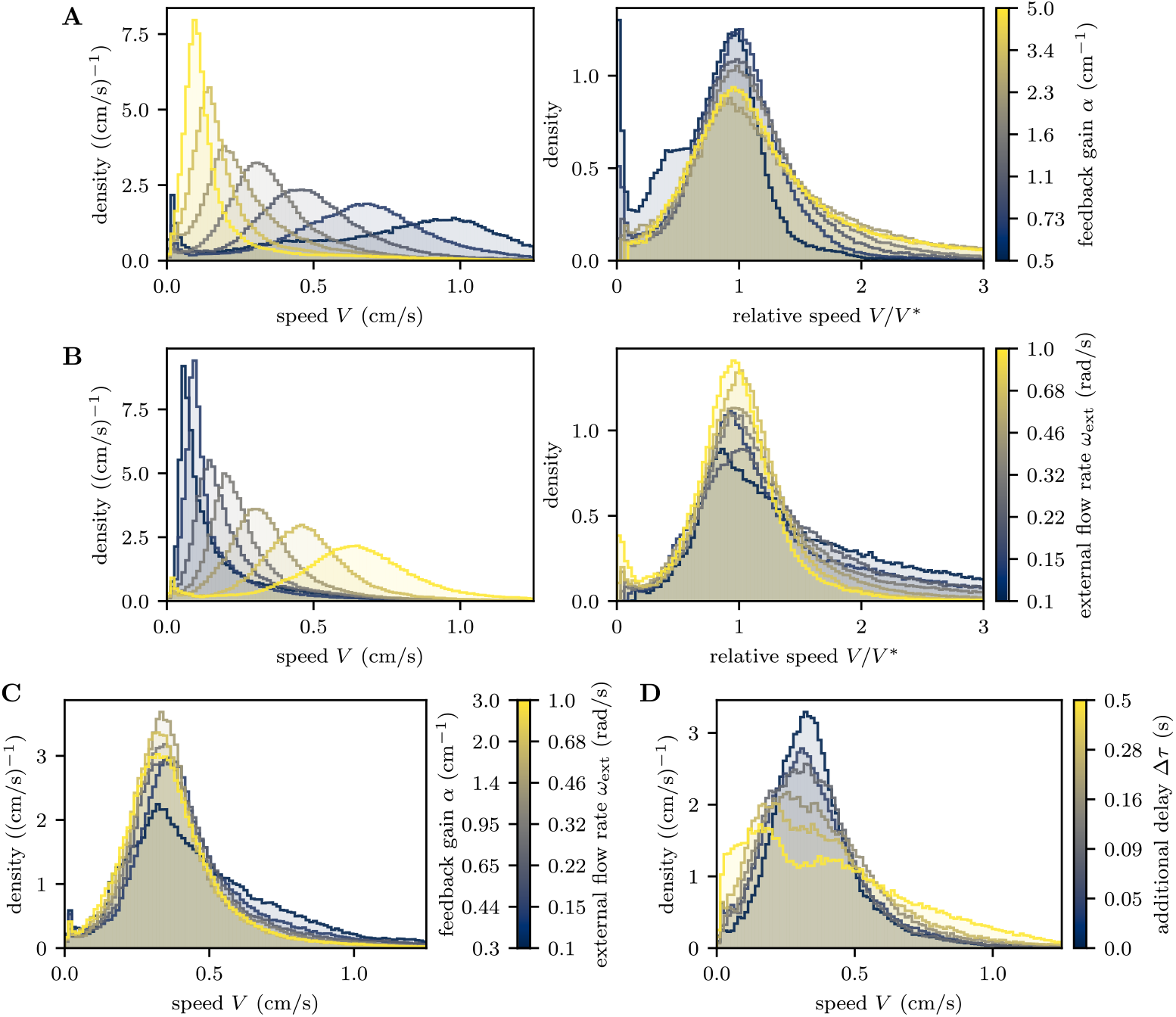
Swimming speed distributions for different conditions. (A) Distributions of swimming speeds (left) pooled across all fish for the experiments with constant currents for different values of *α*. After normalizing the swimming speed by the target speed, all distributions approximately collapse onto a single curve (right). (B) Same as (A), but for the experiments with constant currents for different values of *ω*_ext_. (C) Distributions of swimming speeds (left) pooled across all fish for the experiments with constant currents for different values of *α* and *ω*_ext_ and a fixed *V* ^*\**^. (D) Same as (C), but for the experiments with constant currents for different values of Δ*τ*.

**Figure S6:**
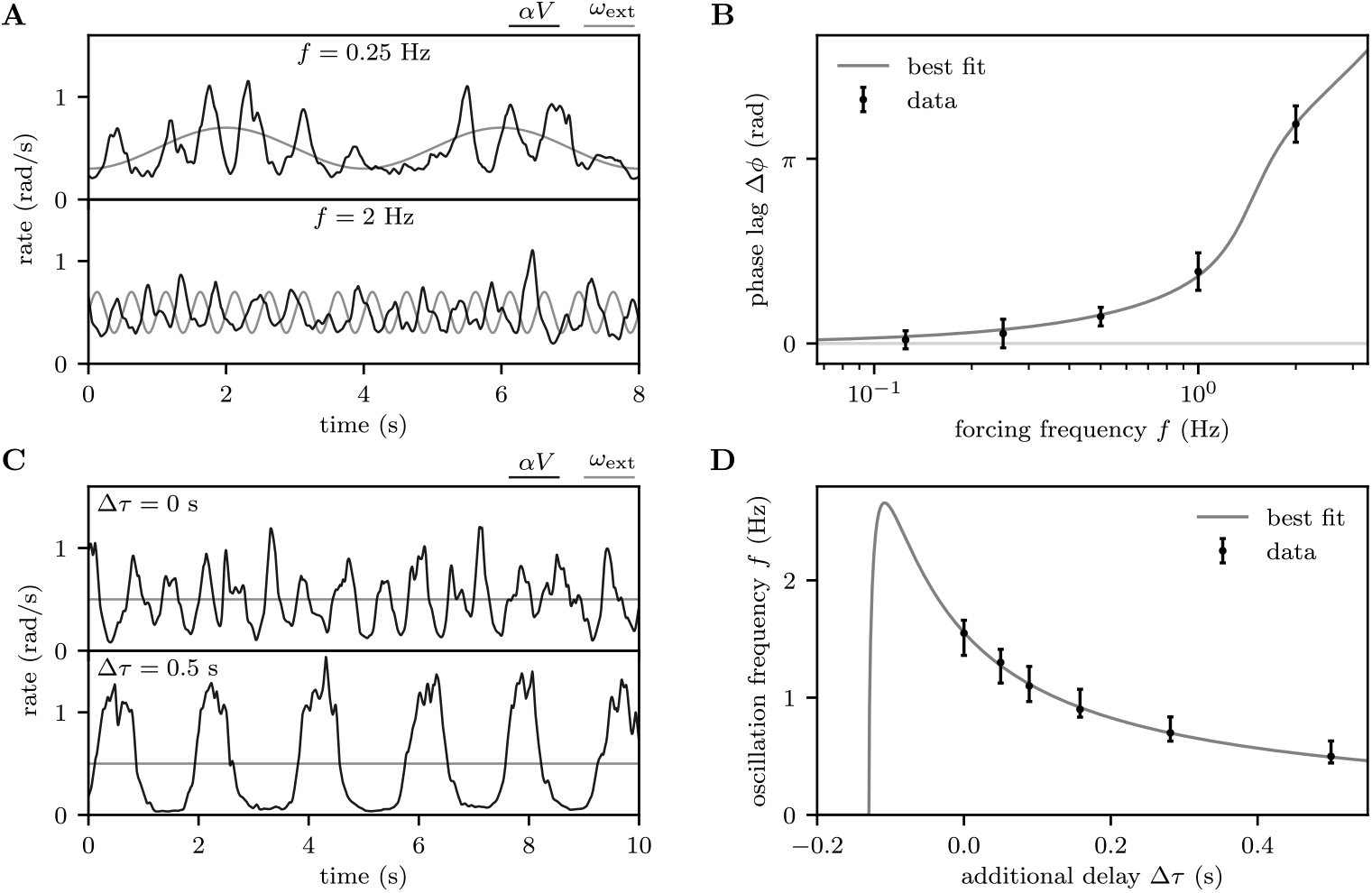
Response to sinusoidal forcing and increased feedback delay. (A) Examples trials showing the response of the fish to currents with sinusoidally varying flow rate for two different frequencies. (B) Phase lag between current speed and fish speed obtained by crosscorrelation for different forcing frequency (black error bars, mean and standard deviation over *N* = 11 fish). The linear delay equation predicts a functional dependence (Eq. 23, gray line) that fits well the data. (C) Examples trials showing the response of the fish to a constant current for different values of the additional feedback delay Δ*τ*. (D) Characteristic frequency of the speed oscillations for different additional feedback delays (black error bars, median of the peak and half-maximum points of the power spectra of 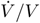 across *N* = 14 fish). The linear delay equation predicts a functional dependence (Eq. 24, gray line) that fits well the data.

**Figure S7:**
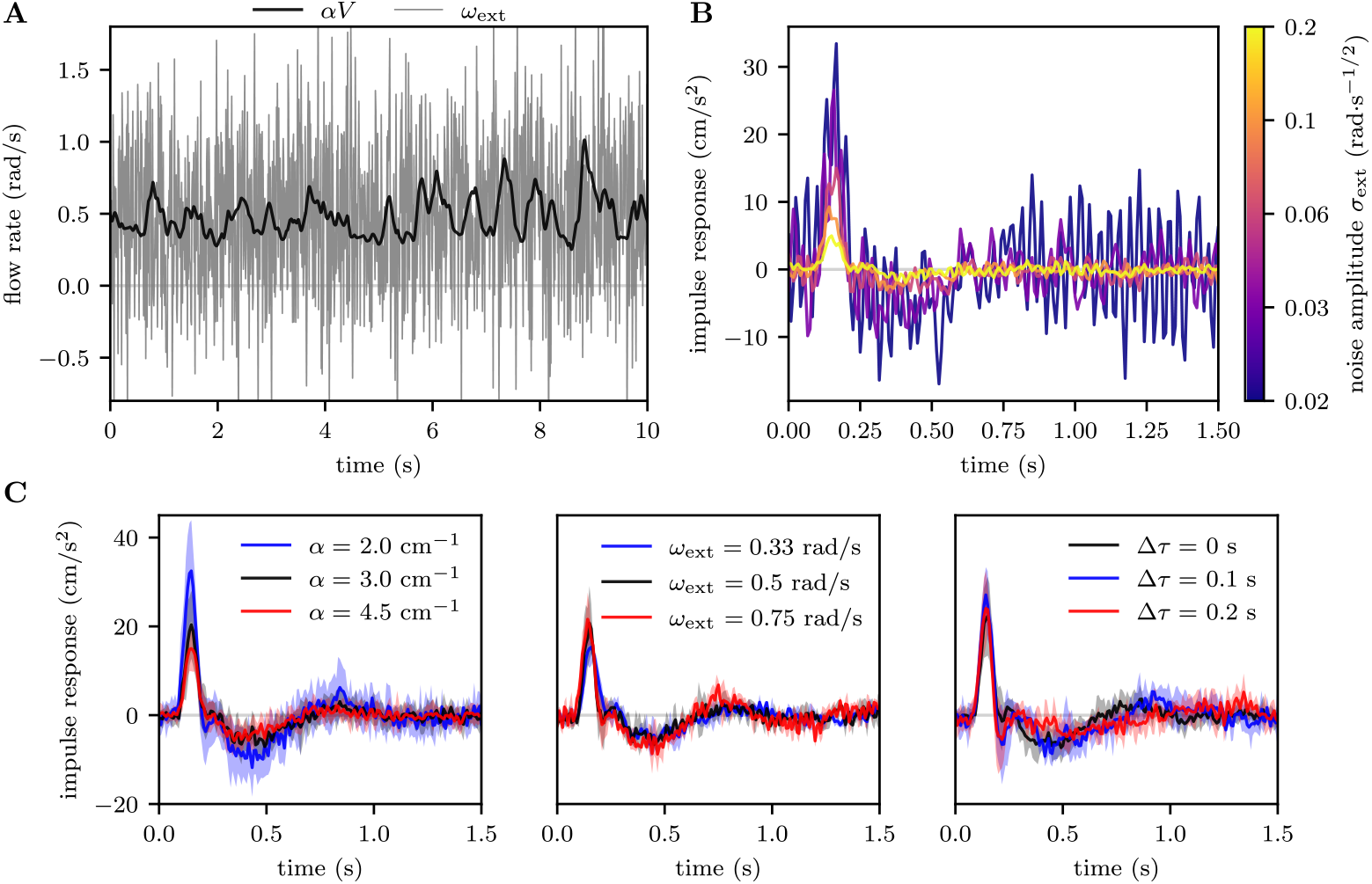
Parameter dependence of the estimated linear impulse response function. (A) Example response of a fish to a current with a flow rate that fluctuates randomly according to a Gaussian distribution. (B) Estimated impulse responses for one example fish for different values of the noise amplitude *σ*_ext_. The shape of the response kernel remains the same but its amplitude peak decreases with *σ*_ext_. This reflects the adaptation of the responsiveness *k*, which is given by its integral. (C) Estimated impulse responses for different values of the feedback gain (left, mean and standard deviation across *N* = 10 fish), external flow rate (middle, *N* = 3 fish), and feedback delay (right, *N* = 6 fish). The shape of the response kernel remains unchanged across all conditions but its amplitude decreases with *α* and increases slightly with *ω*_ext_.

**Figure S8:**
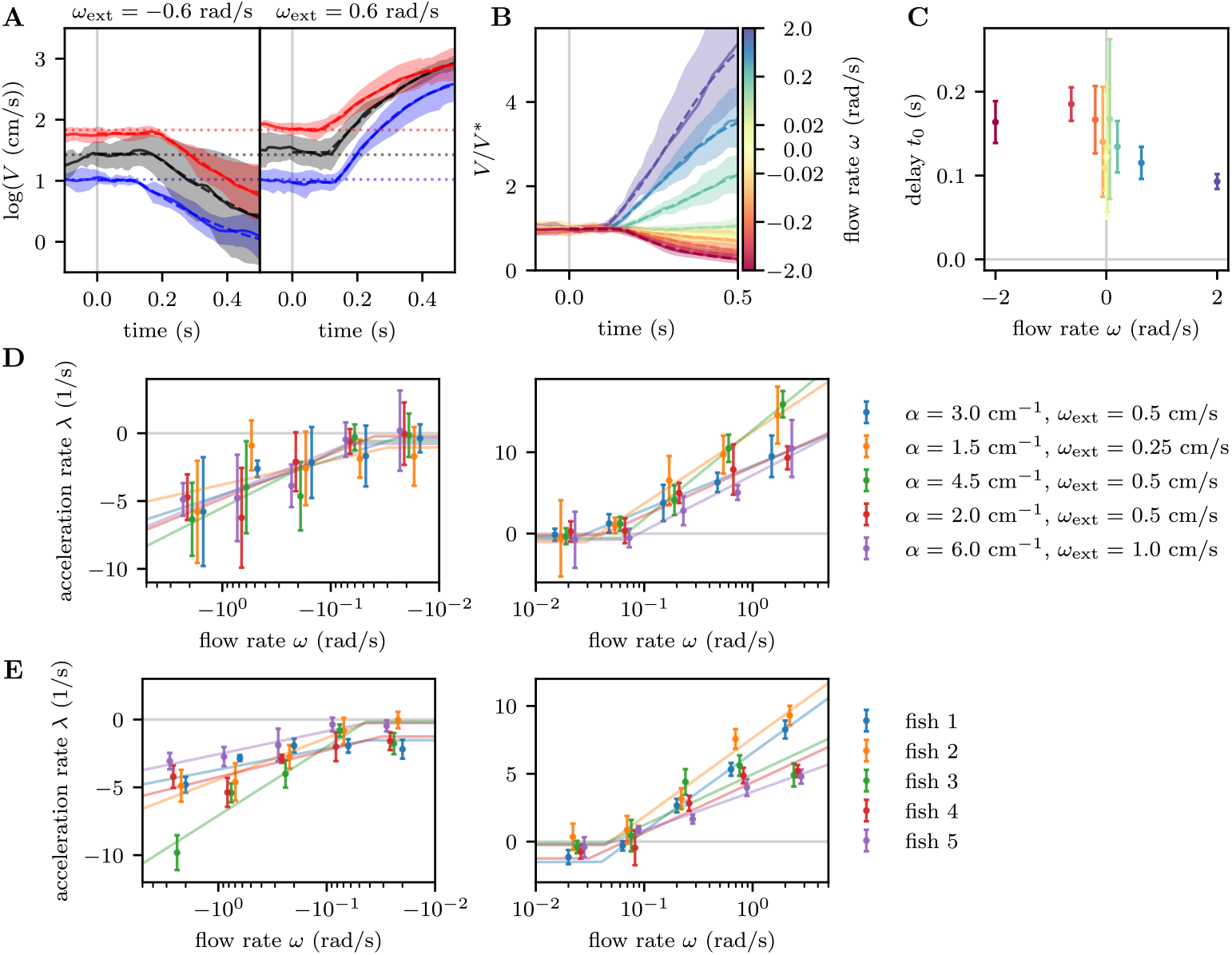
Response to open-loop perturbations. (A) Evolution of the swimming speed (solid lines and shaded areas, mean and standard deviation across *N* = 4 fish) on a logarithmic scale, showing the dependence on the target speed (dotted lines) before the perturbation. (B) Same as Fig. 5A in the main text, but plotted on a linear scale. (C) Delay in the response following the perturbation as a function of the perturbation flow rate (mean and standard deviation across *N* = 5 fish). (D) Acceleration rates for trials with different values of *α* and *ω*_ext_ before the perturbation (see legend, for fish 2 in (E)). The data is shown in semilogarithmic scale for both negative and positive *ω* and it has been slightly shifted horizontally for different conditions to ease visualization. Each trial was fit individually to extract the acceleration rate and the error bars denote the distributions across trials (median and median absolute deviation). The data points are fit with a Eq. 26 for each condition (solid lines). (E) Acceleration rate shown separately for all fish, similarly to (D). The median trajectories across all trials were fit for each fish and the error bars denote the mean and standard deviation of the fit estimates.

**Figure S9:**
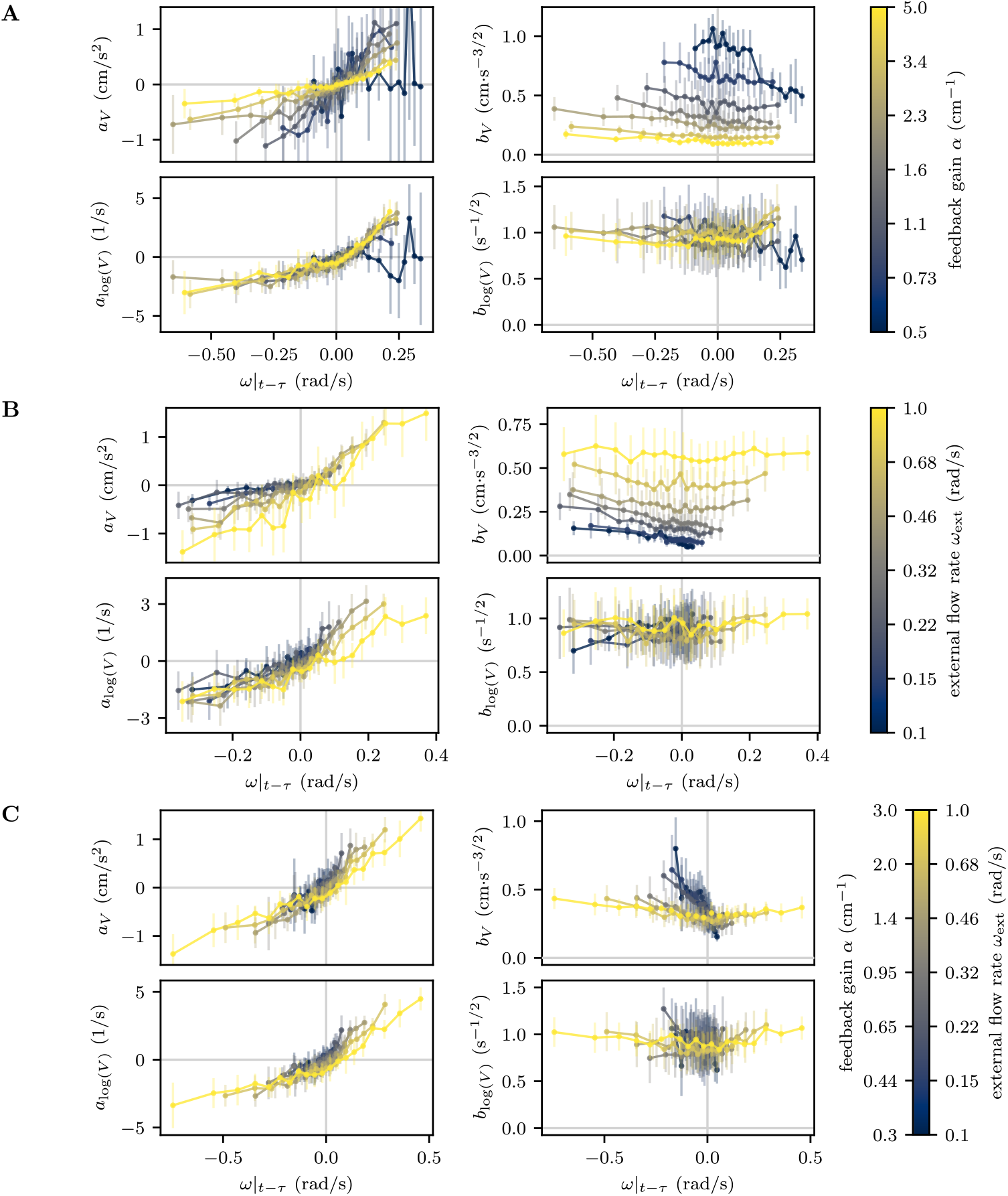
Estimated drift term and noise amplitude for the dynamics of *V* and log(*V*). (A) Drift term and noise amplitude as a function of the perceived flow rate for the experiments with constant currents for different values of *α*. The error bars give the median and median absolute deviation of the estimates evaluated on the data pooled across all fish. The left and right plots show the drift terms *a* and noise amplitudes *b*, respectively. The top plots show the estimates for the dynamics of *V* and the bottom ones for log(*V*). (B) Same as (A), but for different values of *ω*_ext_. (C) Same as (A), but for different values of *α* and *ω*_ext_ and a fixed *V* ^*\**^.

**Figure S10:**
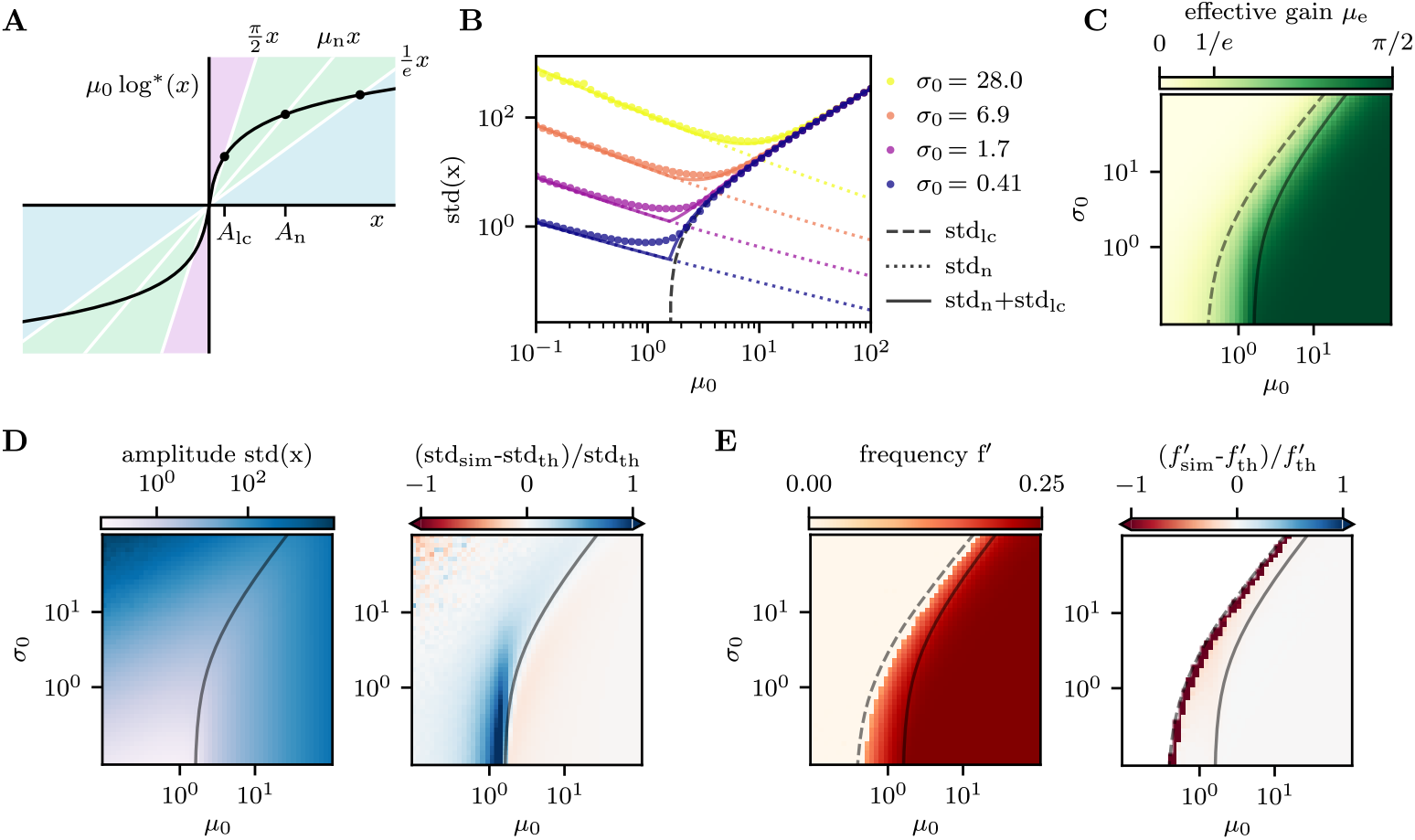
Effective linear theory for the stochastic nonlinear delay equation. (A) Schematic graph showing the definition of the effective amplitudes *A*_lc_ and *A*_n_. (B) Comparison of the fluctuation amplitude computed from simulations of Eq. 34 and from the effective linear theory, for different values of *µ*_0_ and *σ*_0_. The simulations (dots) match the prediction for the limit cycle amplitude (dashed line) for large *µ*_0_ and those for the noise-induced amplitude (dotted lines) for small *µ*_0_. The sum of the two effects (solid lines) matches the simulations except for a small deviation at the transition between the two regimes. (C) Theoretical prediction of the effective gain *µ*_e_. We indicated the curve corresponding to the transition to oscillatory behavior (dashed line, *µ*_e_ = 1*/e*) and the one where the contributions of noise and limit cycle are equal (solid line, std_n_ = std_lc_). (D) Fluctuation amplitude extracted from simulations of Eq. 34 for different values of *µ*_0_ and *σ*_0_ by computing the standard deviation of the sampled *x* values (left) and relative error between the simulations and the theoretical predictions (right). (E) Same as (D), but for the characteristic frequency of the simulated trajectories, extracted from the first local minimum of the autocorrelation.

**Figure S11:**
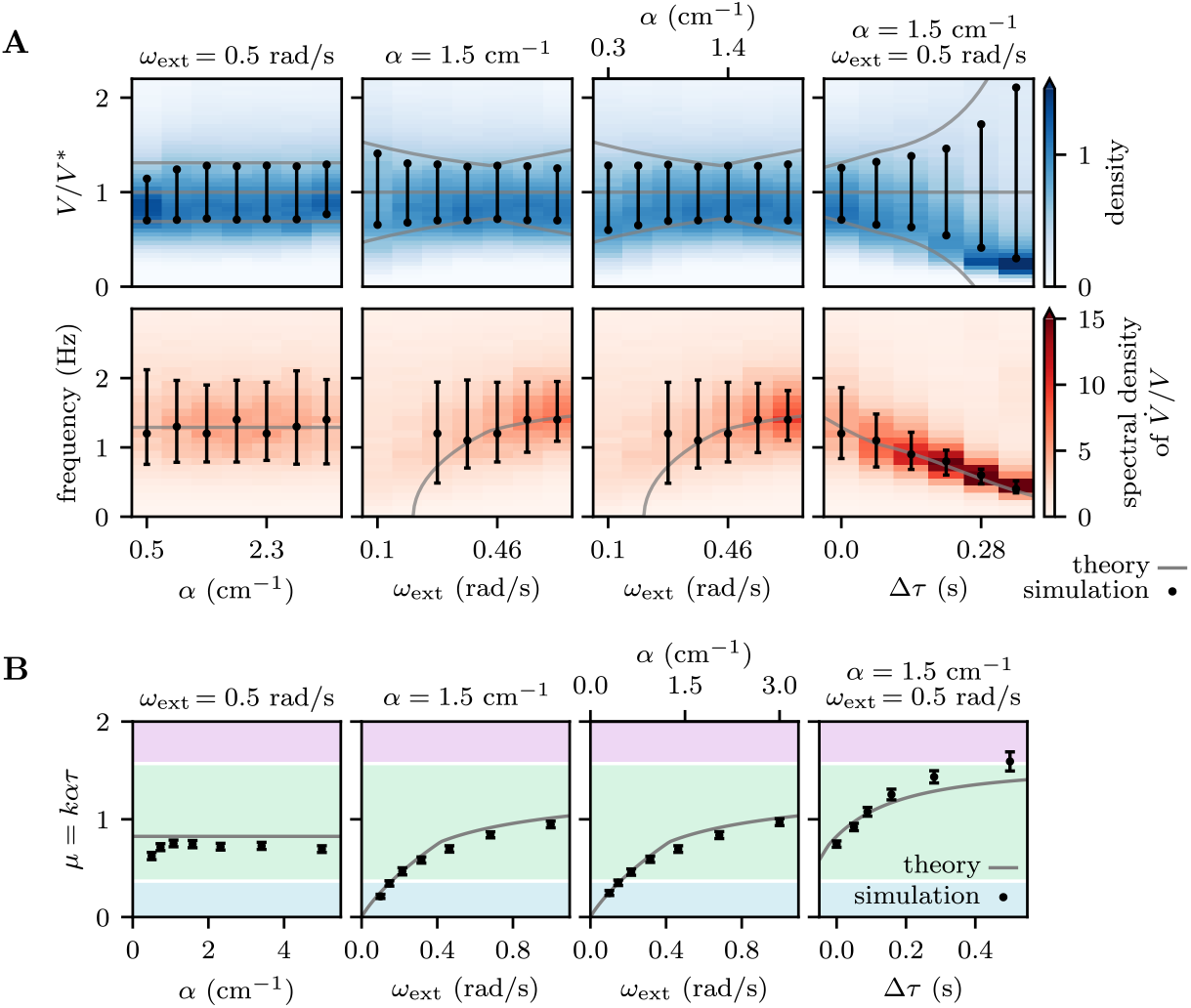
Results of numerical simulations of the model. (A) Distributions of relative swimming speeds *V/V* ^*\**^ (blue, bars indicate the interquartile range) and power spectra of relative acceleration 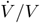 (red, error bars indicate peak and half-maximum values), analogous to Fig. 5E in the main text, but for numerical simulation of Eq. 44. (B) Estimated values of the dimensionless gain *µ* (black error bars, mean and standard deviation of bootstrap samples) by regressing the acceleration 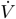 on the optic flow rate *ω* a time *τ* in the past, analogous to Fig. 4B in the main text, but for numerical simulation of Eq. 44.

**Figure S12:**
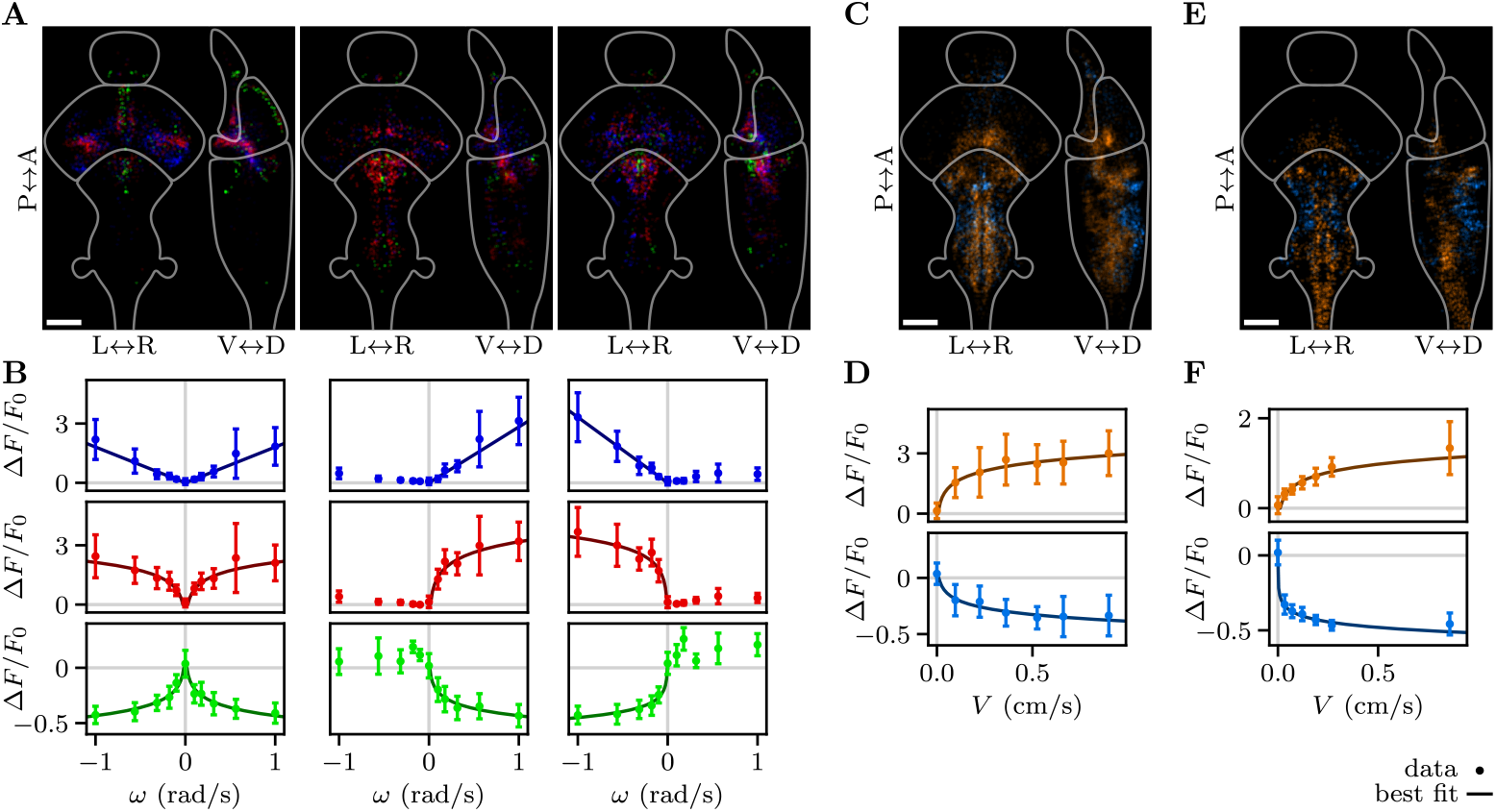
Neuronal populations related to optic flow rate and swimming speed for one example fish. (A-D) Spatial distributions and tuning curves for the neuronal populations correlated with optic flow rate in different directions and with swimming speed for one of the fish used in Fig. 6 in the main text. The figures are analogous to those in the main text. The mean activity of each neuronal population is shown in the tuning curves, where the error bars denote the mean and standard deviation across all the time points corresponding to each bin. (E-F) Spatial distributions and tuning curves for the neuronal populations correlated with swimming speed in one fish during closed-loop stabilization, the analysis is analogous to (C-D).

**Figure S13:**
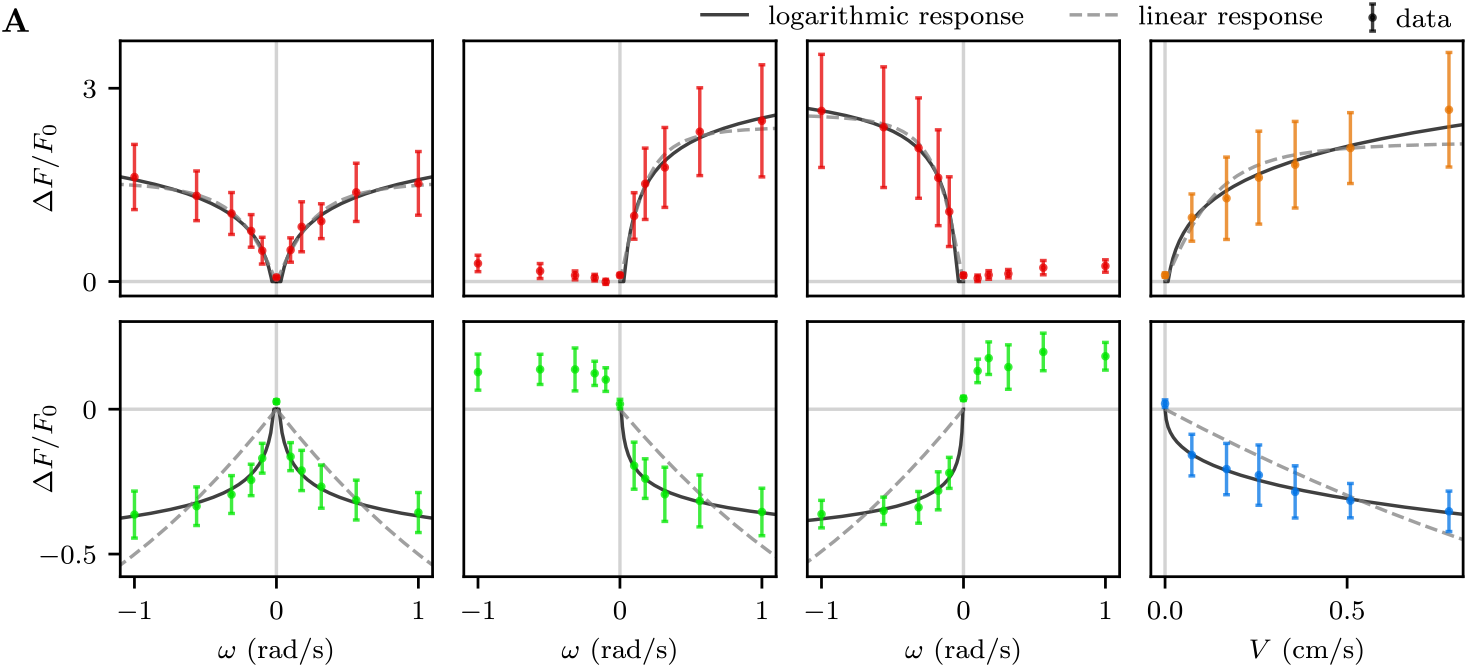
Effect of the calcium indicator saturation on the fits of the tuning curves. (A) Tuning curves for the neuronal populations found to logarithmically encode optic flow rate and swimming speed (same as shown in the main text). The solid and dashed lines are the best fits when taking into account the nonlinearity of the calcium indicator, under the assumption of linear (dashed lines) or logarithmic responses (solid lines).

**Movie S1**. Example of a fish actively turning in the virtual reality to align to external currents oriented in different directions. The movie shows the visual stimulus presented to the fish (top, left), the video recording of the tail with overlaid segmentation (top, middle), a schematic of the fish orientation in the virtual environment, with the fish velocity vector in black and the external current orientation in grey (top, right), and the traces of flow rates and orientations relative to fish and current (bottom). 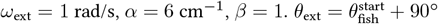 for the first trial and 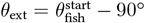 for the second.

**Movie S2**. Example of a fish swimming in the virtual reality and stabilizing its position against forward-directed currents with different flow rates. The movie shows the visual stimulus presented to the fish (top, left), the video-recording of the tail with overlaid segmentation (top, right), and the traces of flow rates relative to fish and current (bottom). *α* = 6 cm^*−*1^, *β* = 1. *ω*_ext_ = 1.5 rad/s for the first trial and *ω*_ext_ = 0.67 rad/s for the second.

**Movie S3**. Example of a fish swimming in the virtual reality to stabilize external currents for different values of the feedback gain. The movie shows the visual stimulus presented to the fish (top, left), the video-recording of the tail with overlaid segmentation (top, right), and the traces of flow rates and speeds relative to fish and current (bottom). *ω*_ext_ = 1 rad/s, *β* = 1. *α* = 9 cm^*−*1^ for the first trial and *α* = 4 cm^*−*1^ for the second.

**Movie S4**. Example of the brain activity of a fish while it stabilizes against external currents in the virtual reality. The movie shows the recorded sequences of 4 different sections of the brain (top) and the traces of flow rates relative to fish and current (bottom). Time is sped up by a factor of 10. *α* = 1.5 cm^*−*1^, *β* = 0.25, *ω*_ext_ = (0.22, 0.33, 0.5) rad/s for the different trials.

**Movie S5**. Visualization of the neuronal populations identified through correlation analysis. The movie shows the three-dimensional spatial distribution of the identified neuronal populations (average densities corresponding to the projections shown in the main text) mapped onto the same reference stack. The densities are normalized so that a single neurons contributes a fraction of 0.1 of the maximum hue intensity. The densities are overlaid on top of the reference stack, shown in grayscale. The movie shows the densities layer by layer moving sequentially from the dorsal side to the ventral side.

**Movie S6**. Visualization comparing the neuronal populations correlated with swimming speed and optic flow in different directions. Analogous to movie S5, but the populations are shown together (independently of positive or negative and linear or logarithmic tuning) to compare the spatial distribution of populations correlated with different signals.

